# Modelling the impacts of imports and non-native subspecies hybridisation in honeybees

**DOI:** 10.1101/2025.07.16.664955

**Authors:** Irene de Carlos, Laura Strachan, Grace P. McCormack, Gregor Gorjanc, Jana Obsteter

## Abstract

Human-mediated movement of organisms for agriculture and ecosystem services often results in hybridisation and introgression between populations of native and non-native species. While introgression may increase genetic diversity, it can erode unique adaptations and reduce fitness, threatening the survival of native lineages. Honeybees offer a good model with extensive records, queen trade and migratory beekeeping facilitating genetic exchange among subspecies.

To explore these dynamics, we used SIMplyBee to simulate hybridisation between populations of native *Apis mellifera mellifera* and non-native *A. m. carnica*. We adopted the parameters from the Irish honeybee population that maintains relatively low levels of import, but is threatened by commercial imports. The model included colony honey yield and fitness as complex polygenic traits. We simulated varying import rates, genetic correlations between fitness in native and non-native environments, and spatial distributions of introgression over 20 years, measuring levels and rate of introgression and genetic means for both traits.

Increased imports accelerated introgression and induced a trade-off between higher honey yield and lower fitness, and decreasing genetic correlations between environments amplified fitness decline. Spatial simulations showed the spread of introgression across the entire simulated area. Halting imports reversed the trend, but purging of introgressed material was slow and varied among replicates. These findings highlight the trade-off between short-term production gains and long-term losses in fitness and adaptation. Our modelling framework provides a reference for exploring introgression in other systems, emphasising that sustainable management of introgression requires restricting imports and breeding locally adapted populations rather than relying on non-native imports.

## 1 Introduction

Hybridisation is the interbreeding of individuals from genetically distinct populations, species, or subspecies. When hybridisation is sustained over generations through back-crossing, it leads to introgression: the integration of genetic material from one lineage into the genetic background of another (Rhymer and Simberloff 1996). At the species level, hybridisation can occasionally result in new species, but it can also result in reduced fertility or inviable offspring (Rhymer and Simberloff 1996). At the subspecies, where reproductive barriers are absent or weak, hybridisation can increase genetic diversity and boost evolutionary potential (Lewontin and Birch 1966). Alternatively, sustained hybridisation and introgression can lead to the loss of unique genetic combinations that have evolved over long periods of adaptation (Rhymer and Simberloff 1996; Benjamin et al. 2021). This process may result in genetic swamping, where native genomes are progressively replaced by admixed genomes, and demographic swamping, where mating with the introduced taxon reduces population growth because hybrids have lower fitness (Allendorf et al. 2001; Rhymer and Simberloff 1996). Hybridisation may also disrupt locally adapted gene complexes, generate outbreeding depression, and reduce the fitness of native populations, especially when rare native taxa are exposed to repeated gene flow from more common introduced populations (Huxel 1999; Bleeker and Martin 2007). Ultimately, these processes can lead to genomic extinction of lineages or subspecies, and homogenise the species (McKinney and Lockwood 1999; De la Rua et al. 2009).

The introduction of non-native species or populations into new environments creates a space for hybridisation. This introduction can be voluntary or involuntary, the latter being a consequence of anthropogenic activity, climate change, or transformation of natural landscapes (Quilodran et al. 2018). These activities have led to a global increase in hybridisation with at least 25% of plant species and 10% of animal species hybridising (Mallet 2005). Agriculture is a major source of voluntary introductions for boosting productivity or providing ecosystem services, but often causing unwanted hybridisation. For example, rainbow trout, widely introduced for hatchery cultiva-tion and sport fishing, has hybridised with local species, driving population declines and extinctions of native species (Metcalf et al. 2008). There are also examples of species introduced for biological pest control hybridising with local populations (e.g. (Havill et al. 2012)). This can have a negative impact through decreased fertility and efficiency, and increased need for chemical control (Corrêa et al. 2019).

Quantifying hybridisation and introgression in wild and managed populations has become increasingly feasible with advances in genomic tools, allowing researchers to estimate admixture levels, identify introgressed genomic regions, and track the timing of hybridisation events (Allendorf et al. 2001). However, empirical estimates alone are insufficient to understand the long-term consequences of introgression or to predict how different management scenarios might affect native populations. Moreover, empirical studies often detect introgression after it has already progressed, at which point management options might be ineffective. For that reason, modelling approaches are becoming increasingly important for anticipating the potential impacts of hybridisation. Computer simulations enable time and cost-effective evaluation of the drivers and long-term consequences of introgression under conditions that are difficult to study empirically and can support proactive management decisions (Quilodran et al. 2018).

Honeybees provide an excellent model for studying hybridisation and its biological consequences, as their lineages, subspecies, and demographic history are well characterised. Moreover, there is active hybridisation since honeybees still live and mate in the wild, and are the subject of anthropogenic activities. In Europe, *Apis mellifera* is structured into three major mitochondrial lineages: Western European (M), Eastern/-Central European (C), and African (A) (Wallberg et al. 2014). Despite this natural differentiation, beekeepers worldwide have long introduced queens across regions, most commonly from the C lineage (De la Rua et al. 2009). In particular, *A. m. ligustica* from Italy and *A. m. carnica* from Central Europe are widespread for their high honey yield, gentle nature, and high fertility (Hoppe et al. 2020; Minozzi et al. 2021; Péntek-Zakar et al. 2015). Beekeepers also transfer whole colonies. In the United States alone, more than two million honeybee colonies are transferred to California every year for crop pollination (Glenny et al. 2017). In Europe, colonies are also transferred to harvest specific types of honey (Martínez-López et al. 2022). These translocations bring together multiple populations at a single location, enabling genetic mixing. Additionally, due to honeybees being managed and economically important, population movements and production data are relatively well documented (Hoppe et al. 2020). However, although there is economic benefit in honeybee trade, imports are contributing to a decline in locally adapted populations.

Introgression has already reduced the natural range of the European dark bee, *A. m. mellifera*, native to north and west Europe, where distinct morphological and behavioural traits could be at risk of being lost (McCann and McCormack 2024; Valentine et al. 2024). This pattern is documented across multiple countries, with introgression reaching 10–30% in several European populations (Pinto et al. 2014; Oleksa et al. 2011; Yanbaev 2024) and continuing to rise with increasing imports (Jones and Semmence 2021; Valentine et al. 2024). Some countries have responded by including *A. m. mellifera* in conservation programmes and banning the import of non-native subspecies into protected areas (Ruottinen et al. 2014).

Despite growing interest in hybridisation and introgression, important knowledge gaps remain. While simulations are widely used in animal and plant genetics (Pook et al. 2020; Gaynor et al. 2021; Bančič et al. 2024), explicit models and applications addressing the drivers and long-term consequences of introgression remain limited. Moreover, the interplay between population parameters, especially the role of GxE interactions, in shaping introgression dynamics and their phenotypic consequences, is poorly understood.

Here, we address these gaps using a stochastic simulation framework applied to honeybee populations. Specifically, we aim to: (i) build a simulation model for assessing long-term hybridisation; (ii) quantify the effect of varying the import percentage on the rate and magnitude of introgression, and key traits, namely production and fitness; (iii) quantify the role of GxE interaction and its interplay with other population parameters; (iv) quantify the speed of introgression spread from a single point of entry; and (v) explore whether populations can recover following a reduction or cessation of imports. To test these hypotheses, we used the population of *Apis mellifera mellifera* in Ireland which represents an excellent model system. In 2018, these populations were reported to be large (Hassett et al. 2018). However, recent increases in honeybee imports coincide with higher observed hybridisation, highlighting the urgent need to assess the effect of imported colonies on both introgression levels and its persistence (Valentine et al. 2024; McCann and McCormack 2024). Furthermore, well-documented records of queen imports are publicly available, facilitating model parameterisation and validation (Department of Agriculture, Food and the Marine 2024). We hypothesised that our model will respond sensibly to changes in import rates reproducing realistic introgression dynamics, that introgression will generate trade-offs between productivity and fitness, and that stronger G×E interactions will exacerbate fitness declines.

## 2 Materials and Methods

The stochastic simulator SIMplyBee (an R package; Obšteter et al. 2023b) was used to simulate a natural yearly cycle of honeybee colonies with import of queens from a non-native population. The R code for this simulation is open-source and available in the supplement and on-line (Data availability: 10). We simulated two populations that resembled a native Irish population of *A. m. mellifera* and a non-native Central European population of *A. m. carnica* (C lineage), each with its own source environment. In both populations, we simulated two complex polygenic traits representing fitness and honey yield. We simulated a genetic correlation in trait expression across the two environments, representing GxE interaction. We simulated 10 years of a burn-in phase followed by 10 years of scenarios varying the percentage of imported queens, the magnitude of GxE interaction for the fitness trait, and the point of entry of the imports. We compared the scenarios based on the level of introgression and the genetic mean for fitness and honey yield.

### 2.1 Simulation

We generated the founder genomes with the Markovian Coalescent Simulator implemented in SIMplyBee, according to *A. mellifera* demographic model (Wallberg et al. 2014; Obšteter et al. 2023b), mutation rate of 3.4 x 10*^−^*^9^ per bp (Beye et al. 2006) and recombination rate of 2.3 x 10*^−^*^7^ per bp (Yang et al. 2015). We generated 500 *A. m. mellifera* and 500 *A. m. carnica* diploid founder genomes. Each genome contained 16 chromosomes with 1000 segregation sites per chromosome. From the founder genomes, we created the base populations of 500 *A. m. mellifera* colonies that represented the population in Ireland, and 500 *A. m. carnica* colonies that represented the population of in Central Europe providing the source of imports of C lineage queens. We generated 10 replicates of these base populations. Each colony was simulated with one queen and 10 workers, which, to reduce computing time, represented the full size of a colony. The analysis itself did not require the simulation of workers, but we simulated a small number to avoid any simulation errors.

Within each native environment, we simulated two complex polygenic traits, fitness and honey yield (Guichard et al. 2023a). We simulated both traits as effected by 100 additive quantitative trait loci (QTLs) per chromosome (1600 QTLs per trait in total) and a trait mean of 0. For honey yield, we set genetic variance to 0.25, and residual variance to 0.75, which resulted in a heritability of 0.25, as estimated for natural honeybee populations (Maucourt et al. 2020). For fitness, we set the genetic variance to 0.1 and residual variance to 0.9, resulting in a heritability of 0.1, typical of fitness-related traits (Kruuk et al. 2000; Guichard et al. 2023b). We modelled the GxE interaction by simulating fitness and honey yield in both environments as genetically correlated traits, resulting in four simulated traits: honey yield in Ireland, honey yield in Central Europe, fitness in Ireland, and fitness in Central Europe. Therefore, each simulated individual had a genetic and phenotypic value for each of the four traits, regardless of their physical location. However, only the “local” values were used for selection, breeding, or import decisions. For honey yield, we fixed the genetic correlation between environments to 0.75, following Hatjina et al. (2014) that showed that honeybees produced more honey in their native environment than in a non-native one. For fitness, we varied the genetic correlation between the scenarios as described in the following.

### 2.2 Burn-in

During the burn-in phase, we kept the two populations closed and simulated 10 years of a beekeeping yearly cycle within each population. This cycle simulated three bee-keeping seasons: spring, summer, and winter, with colony events occurring at different probabilities depending on the season (Table 1). These probabilities were estimated from the current literature (Mattila and Otis 2006; Seitz et al. 2015; Rangel et al. 2013; Brodschneider et al. 2018; Gray et al. 2023). We ran the 15 replicates of the base populations through the burn-in phase.

**Table 1.** Probabilities of colony swarming, supersedure, and collapse in spring, summer, and winter simulated periods.

|  | Spring | Summer | Winter |
| --- | --- | --- | --- |
| Supersede | 0.05 | 0.05 | – |
| Swarm | 0.05 | 0.01 | – |
| Collapse | 0.10 | 0.10 | 0.18 |

In the spring period (Figure 1 A), we first simulated the growth of existing colonies to their full size. Next, we split all colonies to simulate swarming prevention, which generated new colonies that were re-queened with age 0 virgin queens obtained from the original colonies. Then, we swarmed or superseded a percentage of colonies (Table 1). The remnant colonies of the swarms or supersedures were naturally re-queened with their own virgin queens. Next, we generated drones from colonies with queens of age 1 or older to create a drone congregation area (DCA) and mated all the new virgin colonies with an average of 15 random drones from the DCA. Finally, we ended the spring period with the collapse of a proportion of colonies, mimicking queen loss during a nuptial flight (Table 1).

**Fig. 1.**
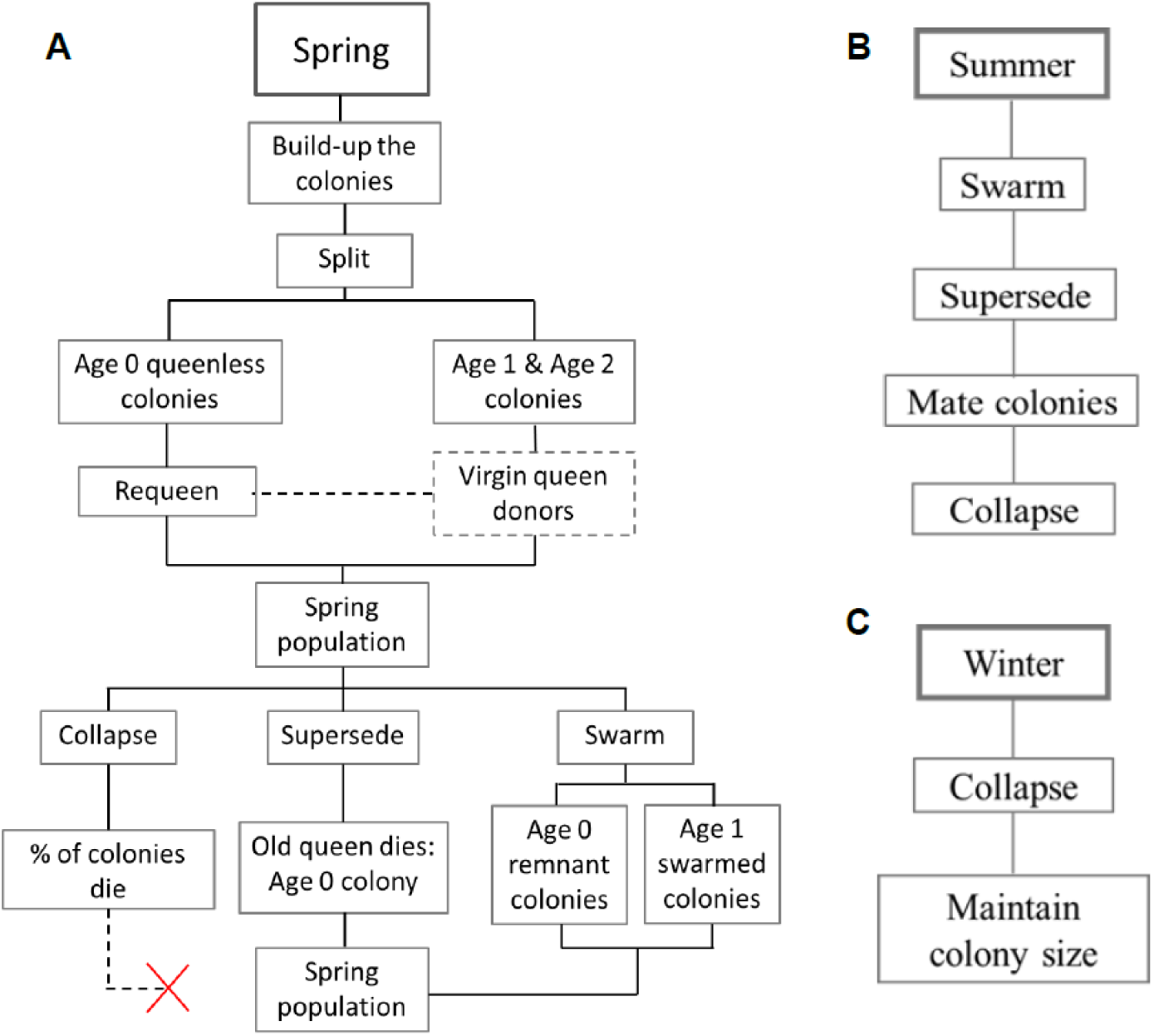
Flow diagram of the simulated yearly cycle of honeybee colonies in A) the spring period, B) the summer period, and C) the winter period

In the summer period (Figure 1 B), we again swarmed a proportion of colonies (Table 1), though less than in the spring period, since swarms occur mostly in spring, when the colonies outgrow their hive (Thomas 2016). We followed this by superseding or collapsing a percentage of colonies (Table 1). The resulting virgin colonies mated as in the spring period.

Finally, in the winter period (Figure 1 C), we collapsed a larger percentage of colonies due to overwintering challenges (Table 1). We collapsed colonies with the lowest phenotypic values for fitness in the corresponding environment to simulate *directional* selection, driving genetic differentiation between the two populations evolving in two different environments. As a result, the Irish population had a higher genetic mean for “fitness in Ireland” than the Central European population at the end of the burn-in phase. At the end of each yearly cycle, we removed some splits to maintain constant population sizes. In the Central European population, we removed the colonies with the lowest phenotypic value for honey yield to simulate directional selection on this trait. This selection reflected the long-term selective breeding for this trait and the presumed superior honey yields of the imported bees into Ireland.

### 2.3 Scenarios

After the burn-in, we continued running the 15 replicates for each of the three sets of scenarios for 10 additional years. In this phase, we transitioned from directional to stabilising selection on fitness, assuming that both populations had reached their optimal fitness levels after the burn-in. To implement this, we selectively collapsed colonies whose phenotypic value for fitness deviated most from the population’s mean fitness during the winter period.

First, we simulated the import of queens from the C lineage into the Irish population by requeening a percentage of spring splits with imported virgin queens (Figure 2). We compared the scenarios with 0% (Control), 1.5%, 4%, and 10% of imports, which were determined based on the reported imports to Ireland (McCann and McCormack 2024).

**Fig. 2.**
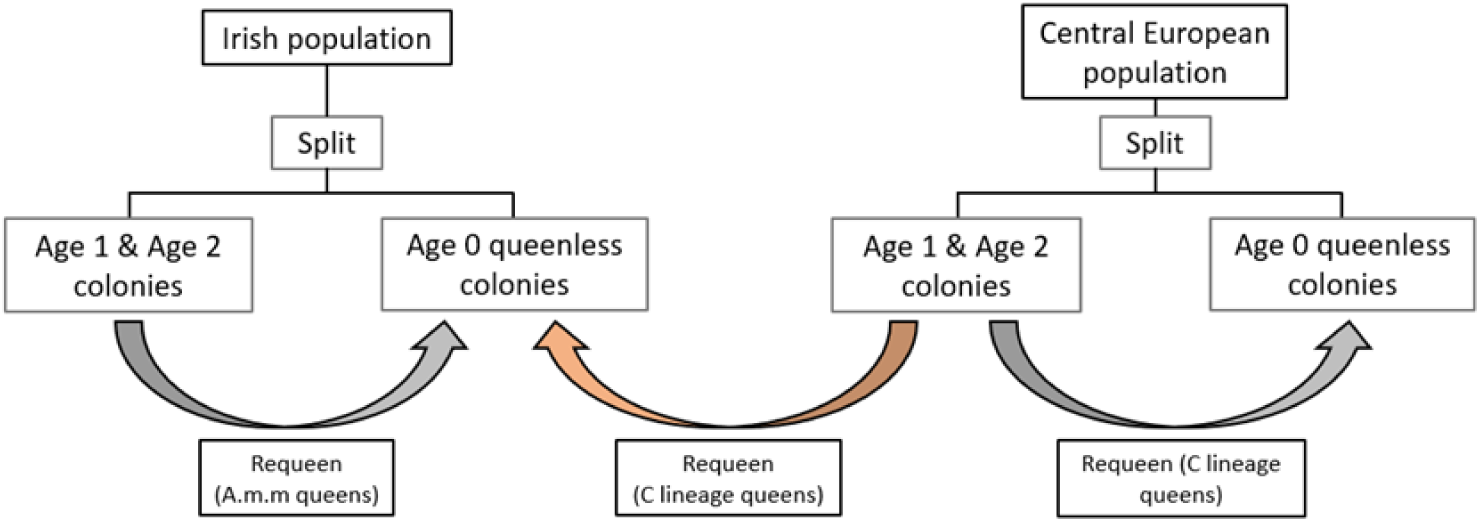
Flow diagram of the import process: re-queening of age 0 queen-less colonies in the Irish population with C lineage queens from the Central European population after the burn-in phase

Second, to explore the effects of adaptation to the local environment and GxE interactions, we created a set of scenarios that varied the genetic correlation between fitness in Ireland and Central Europe. Büchler et al. (2014) found higher survival of honeybees with native genotypes over non-native genotypes, indicating the presence of GxE interactions and a past adaptation to the environment. However, given the lack of studies on the extent of the GxE interaction in honeybees, we evaluated a wide range of genetic correlations: −0.75, −0.25, 0.25, and 0.75 (base scenario).

Third, to explore the effect of the spatial distribution of imports, we created spatially-aware scenarios. In the scenarios described above, the colonies have no notion of the location and can hence mate randomly (panmictic simulation). This was done to simplify the simulations, as we were mostly interested in the qualitative differences between scenarios. However, colony locations and the locations of imports are likely to impact the introgression. SIMplyBee enables spatially-aware simulations, where each colony is assigned a location and the mating is restricted to a user-defined radius. To evaluate the effect of the point of import on the spread of introgression over time and place, we created a scenario by simulating an area with a single point of import at the center of the simulated area. We chose to place the point of import at the center of the simulated area to measure the effects of introgression symmetrically in all directions. We simulated an area of 150*km*^2^ with 500 colonies randomly placed. This mimicked the colony density in county Dublin (Department of Agriculture, Food and the Marine 2019), which receives the majority of imports in Ireland (about 3,000 reported honeybee colonies within 900*km*^2^ in 2020).

The 500 colonies were randomly placed into groups of between 5 and 15 colonies, representing the Irish apiaries. We restricted the mating to 15*km* radius and the new location of the swarms to 1.5*km* radius (Jensen et al. 2005; Schultz et al. 2008). Colonies created as splits were given the same location as the colony they originated from. Virgin colonies were re-queened with virgin queens sampled from a nearby area. For this, we divided the area into 16 smaller areas and sampled one colony as the virgin queen donor for each area.

Lastly, we simulated the effect of halting the import to evaluate if a population can recover after long periods of continuous imports. For this, we extended the spatially-aware scenario with a single point of entry for an additional 10 years (30 simulated years in total), where we halted imports to evaluate if the population can recover after long periods of continuous imports.

### 2.4 Analysis

To assess the level of introgression, we tracked recombination of genomes throughout the simulation and, with it, identity by descent inheritance of alleles from both base populations. This enabled quantifying the proportion of alleles that originated from the native *A. m. mellifera* or the non-native C lineage population. We measured the level of introgression for each queen as 1 minus the proportion of alleles that originate from the founder *A. m. mellifera* alleles. Then we averaged the level over all queens by the year of birth.

We computed the rate of introgression as the slope of a linear model fitting the mean level of introgression over years for each replicate: *lm*(Introgression ~ Year − 1). We omitted the intercept, because the level of introgression was 0 at the end of burn-in. We computed the average slope across the replicates of the scenarios, as well as the 95% quantiles.

We computed the genetic trend for “honey yield in Ireland” and “fitness in Ireland” traits by calculating the mean genetic value of queens by their year of birth in the Irish and the Central European populations for each replicate. These values were centered to have a mean of 0 in the last year of burn-in, year 10. Then we computed the average across the replicates of the scenarios as well as the 95% quantiles.

To assess the contribution of imported queens to genetic gain, we next partitioned the overall genetic trends for honey yield and fitness traits in the Irish population to the contribution of native *A. m. mellifera* and non-native C lineage population with AlphaPart R package (Obšteter et al. 2021).

In the spatially-aware scenario, we assessed the spread of introgression by plotting the proportion of introgression against the Euclidean distance of the colonies from the point of import. The Euclidean distance *d* between a colony located at (*x_i_, y_i_*) and the point of entry at (75, 75) (centre of the 150*km*^2^ area) was calculated as 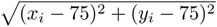.

Moreover, we analysed the cline of introgression, to study gradual change across geographical distance. We fitted clines for each year of importation using the Hybrid Zone Analysis R package (Derryberry et al. 2014). For each year of import, we collected the estimated c value, representing the centre of the cline. We averaged the c values across replicates to obtain the mean c value per year, as well as the 95% quantiles.

We performed ANOVA to assess the significance of differences in the collected quantities. We performed the post-hoc Tukey HSD tests to evaluate the differences between levels of a specific factor. All of these analyses were performed using the base R packages (R Core Team 2024).

## 3 Results

The results show (1) the effects of varying percentage of imports and the genetic correlation between fitness in two environments on the level and rate of introgression and mean honey yield in the Irish population, (2) the dispersion of introgression with a single point of entry, and (3) effect of halting imports. Increasing imports resulted in a proportional increase in introgression and honey yield; however, it also resulted in a notable decrease in fitness. Changing the genetic correlation between fitness in two environments did not significantly affect introgression or honey yield, but negative correlations led to a stronger decrease in fitness. Introgression from a single point of entry spread fast through the simulated area over the years. Halting imports led to a decay in introgression levels, but the rate of this decay was low.

### 3.1 Percentage of imports

The results showed that the level and rate of introgression increased from year 10 to year 20 in all scenarios that imported non-native queens (Figure 3). The increase became significantly more evident with higher import percentages (Supplementary information: Additional file 1 and Additional file 2). We observed the highest rate of introgression in the scenario with 10% imports (0.013). Here, the level of introgression reached 0.39 by year 20, meaning that 39% of the population’s genome was non-native. Decreasing the percentage of imports decreased the level of introgression, with the population being 18.3% and 7.4% introgressed in year 20 respectively with 4% and 1.5% imports (Table 2). The control scenario had no import and hence retained the level and rate of introgression at 0.

**Fig. 3.**
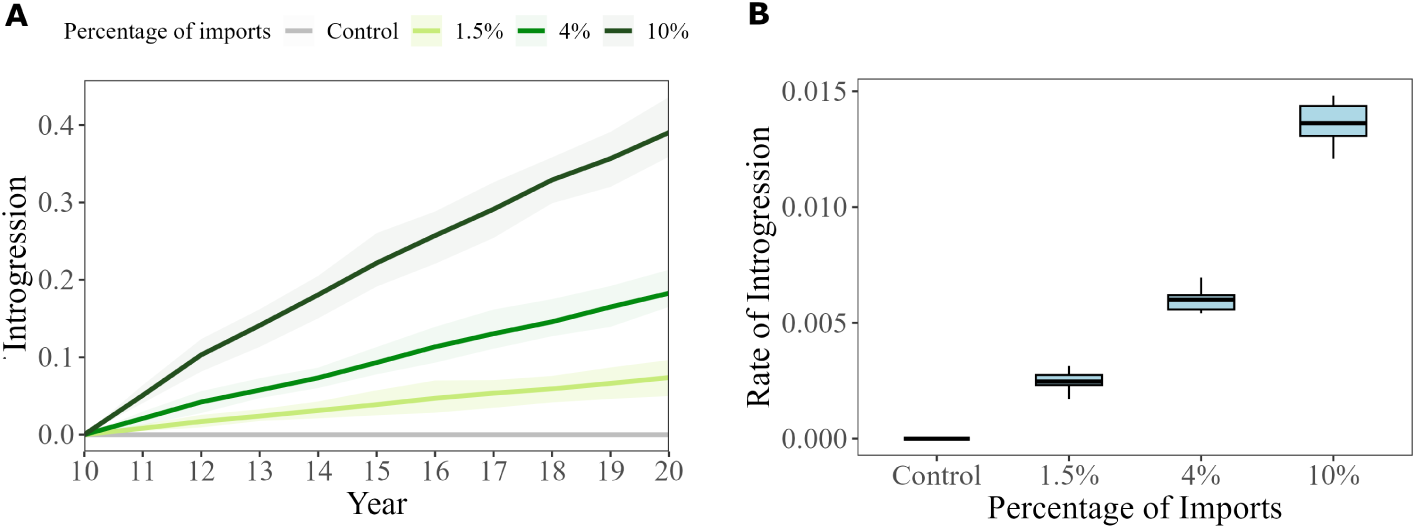
Changes in the level and rate of introgression across replicates in scenarios with the varying percentage of imports of the C lineage into the Irish population: Control (0%), 1.5%, 4%, and 10%. A) The trend in mean level of introgression with the corresponding shaded regions indicating the 95% quantiles. B) Distribution of introgression rates across replicates.

**Table 2.**
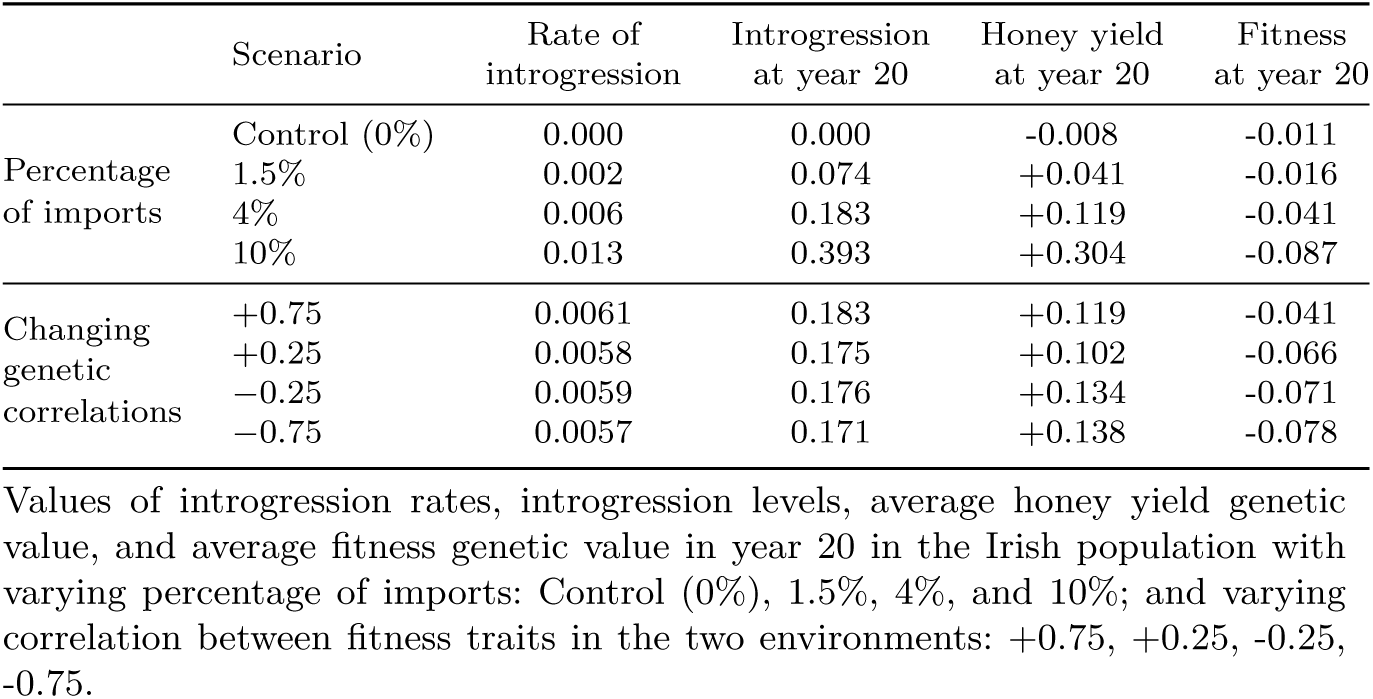
Introgression metrics and genetic values in year 20 under different import and correlation scenarios.

|  | Scenario | Rate of introgression | Introgression at year 20 | Honey yield at year 20 | Fitness at year 20 |
| --- | --- | --- | --- | --- | --- |
| Percentage of imports | Control (0%) | 0.000 | 0.000 | -0.008 | -0.011 |
|  | 1.5% | 0.002 | 0.074 | +0.041 | -0.016 |
|  | 4% | 0.006 | 0.183 | +0.119 | -0.041 |
|  | 10% | 0.013 | 0.393 | +0.304 | -0.087 |
| Changing genetic correlations | +0.75 | 0.0061 | 0.183 | +0.119 | -0.041 |
|  | +0.25 | 0.0058 | 0.175 | +0.102 | -0.066 |
|  | -0.25 | 0.0059 | 0.176 | +0.134 | -0.071 |
|  | -0.75 | 0.0057 | 0.171 | +0.138 | -0.078 |

Values of introgression rates, introgression levels, average honey yield genetic value, and average fitness genetic value in year 20 in the Irish population with varying percentage of imports: Control (0%), 1.5%, 4%, and 10%; and varying correlation between fitness traits in the two environments: +0.75, +0.25, -0.25, -0.75.

In accordance with the introgression, the mean genetic value for honey yield in the Irish population increased with increasing percentage of imports, while the mean genetic value for fitness decreased. In the control scenario, traits fluctuated around zero as honey yield in Ireland was not under directional selection, fitness was under stabilising selection at the optima, and there were no imports (Figure 4). We observed the highest increase in mean genetic value for honey yield in the scenario with 10% import, reaching +0.304 genetic standard deviations by year 20. This value was about 7 times higher than in the 1.5% import scenario and about 3 times higher than in the 4% scenario (Table 2, Supplementary information: Additional file 1). However, the 10% import scenario showed a significant and steep decrease in mean genetic value for fitness, dropping to −0.087 genetic standard deviations by year 20, while the decrease in fitness in the 4% and 1.5% scenarios were less pronounced and not significantly different from the control group (Supplementary information: Additional file 2).

**Fig. 4.**
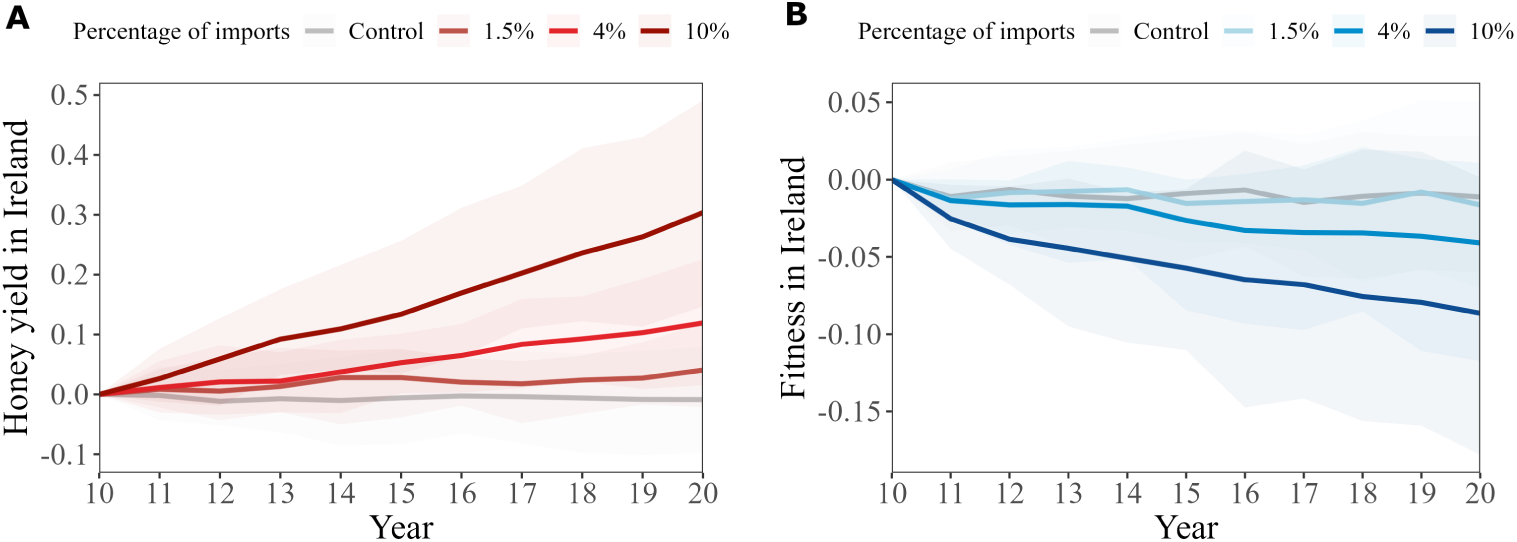
Changes in the mean genetic value for (A) honey yield (red lines) and (B) fitness trait (blue lines) in the Irish population from year 10 to year 20 in scenarios with a varying percentages of imports: Control (0%), 1.5%, 4%, and 10%. The solid lines represent the mean genetic value across replicates with corresponding shaded regions indicating the 95% quantiles.

The partition of genetic trends for both traits further confirmed the positive contribution of imported queens to honey yield, with the imports accounting for 99% of the increase. This partition also confirmed the negative contribution of imported queens to fitness, with the imports accounting for 99% of the decrease (Supplementary information: Additional file 3 and Additional file 4).

### 3.2 Genetic correlation between fitness in the two environments

The results revealed that the genetic correlation between fitness in Ireland and Central Europe had limited effects on the level and rate of introgression (Figure 5). Although the differences in introgression matched our expectations, there was a substantial variation between simulation replicates and the differences between scenarios were not statistically significant (Supplementary information: Additional file 1 and Additional file 2). The high genetic correlation of 0.75 gave the highest rate of 0.0061/year and the level of 0.183 in year 20. The low genetic correlation of −0.75 gave the lowest rate of 0.0057/year and the level of 0.171 in year 20.

**Fig. 5.**
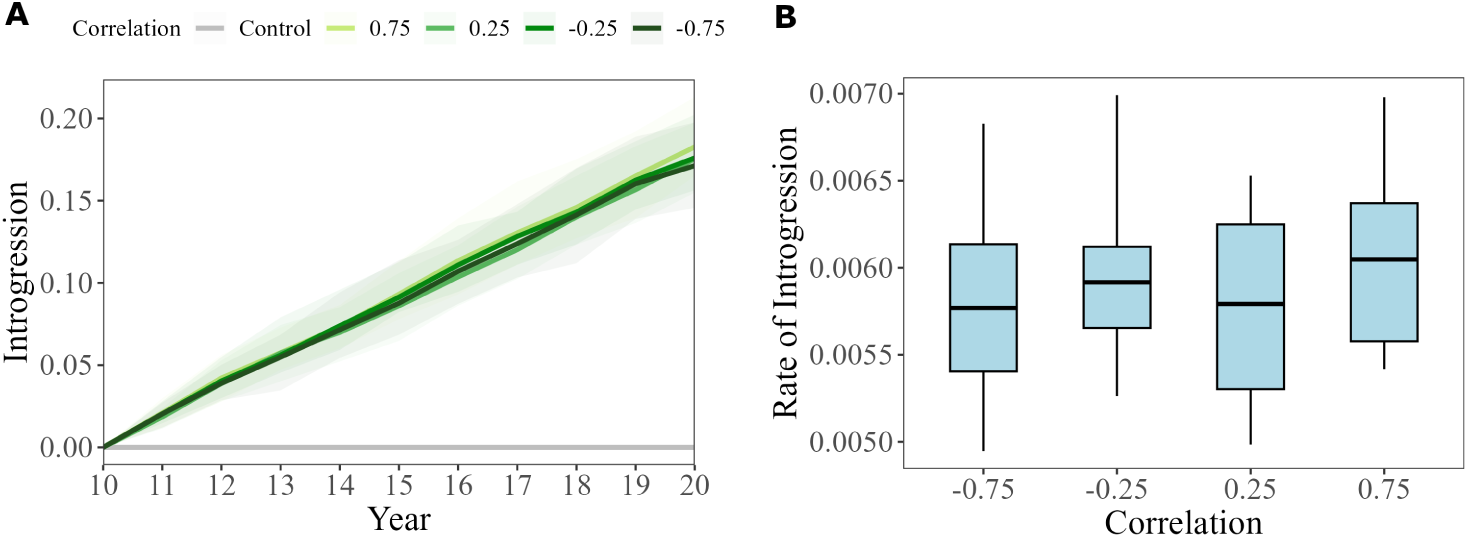
Changes in the level and rate of introgression from year 10 to year 20 across replicates in scenarios with a varying genetic correlations between fitness traits in Ireland and Central Europe: +0.75, +0.25, −0.25, −0.75. The percentage of imports was set to 4% in all scenarios. A control population that received no imports is also represented. A) Trend in the introgression level with corresponding shaded regions indicating the 95% quantiles. B) Boxplot of the rate of introgression.

The mean genetic value for honey yield in the Irish population by year 20 did not follow a pattern regarding genetic correlation for fitness. With the −0.75 correlation scenario exhibiting the highest mean (0.138 genetic standard deviations) and the 0.25 scenario exhibiting the lowest (0.102 genetic standard deviations) (Figure 6, Table 2). However, none of the differences between the scenarios were significant given the high variation between replicates (Supplementary information: Additional file 1 and Additional file 2).

**Fig. 6.**
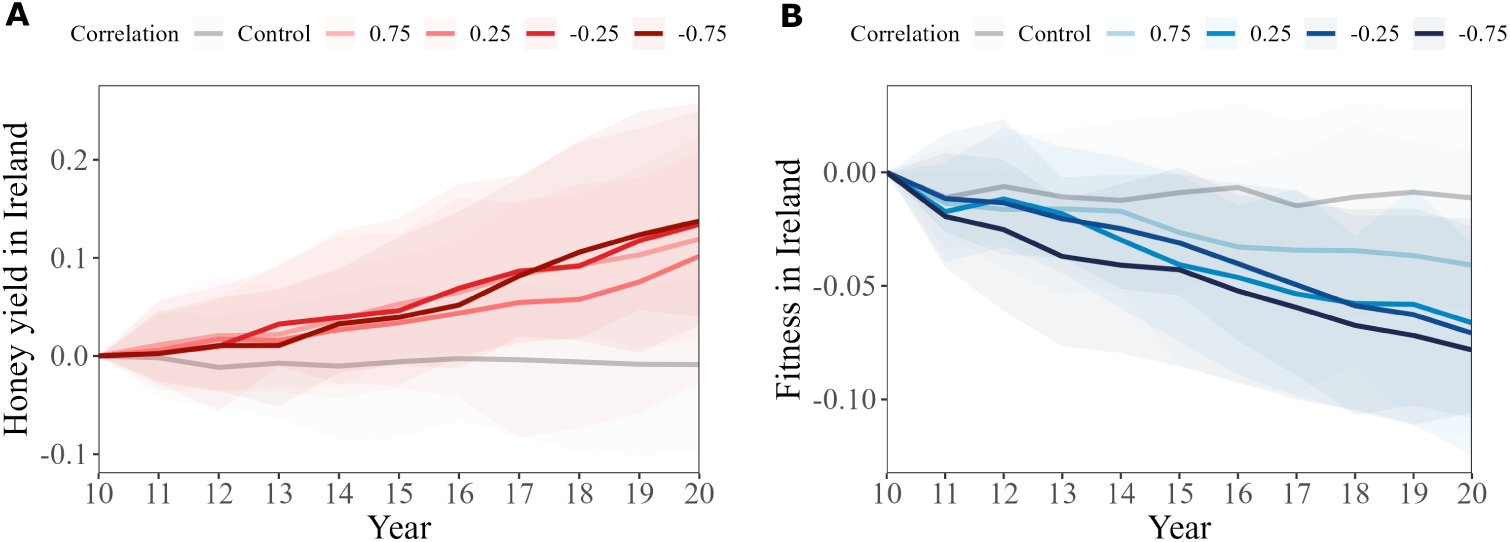
Changes in the mean genetic value for (A) honey yield (red lines) and (B) fitness trait (blue lines) in the Irish population from year 10 to year 20 in scenarios across varying correlations between fitness traits: +0.75, +0.25, −0.25, −0.75. The solid lines represent the mean genetic value across replicates with corresponding shaded regions indicating the 95% quantiles. A control population that received no imports is also represented.

The partitioning of genetic trends for honey yield further confirmed that although the import increased the honey yield in all scenarios, there was no clear pattern in the size of contribution related to the genetic correlation for fitness in the two environments (Supplementary information: Additional file 5).

The mean genetic value for the fitness trait in the Irish population by year 20 decreased with decreasing the genetic correlation between fitness trait in Ireland and Central Europe (Figure 6, Supplementary information: Additional file 1). The scenario with the +0.75 exhibited the smallest decrease in the value of fitness by year 20 (−0.041 genetic standard deviations), while the scenario with the −0.75 correlation exhibited the most significant decrease in the value of fitness by year 20 (−0.078 genetic standard deviations) (Table 2, Supplementary information: Additional file 2).

The partition of genetic trends for fitness further revealed the negative contribution of the imported honeybees to the genetic trend for fitness in the Irish population. While the domestic contribution to fitness fluctuated around zero due to stabilising selection, the contribution of the imports was increasingly negative as we decreased the fitness genetic correlation, with the maximum decrease observed with the −0.75 genetic correlation. Fitness declines were mostly driven by the imports, which accounted for 99% of the observed decline (Supplementary information: Additional file 6)

### 3.3 Point of entry

In the scenario with a single point of entry, colonies closer to the entry point exhibited higher levels of introgression compared to those further away. For instance, colonies located in a radius of 25km or closer to the point of entry showed levels of introgression that ranged between 75% to 100%, while colonies 100km away from the entry point showed no introgression over the simulated period (Figure 7a). The extensive spread of C lineage alleles was evidenced by the presence of some level of introgression even in colonies located 80km away from the point of entry at year 20. Moreover, the cline analysis revealed a significant increase in the *c* value over the years (p < 0.001) (Supplementary information: Additional file 8). Hence, the centre of the cline was moving away from the point of entry, also indicating that introgression was spreading progressively further from the point of entry as years passed (Figure 7b, Supplementary information: Additional file 7).

**Fig. 7.**
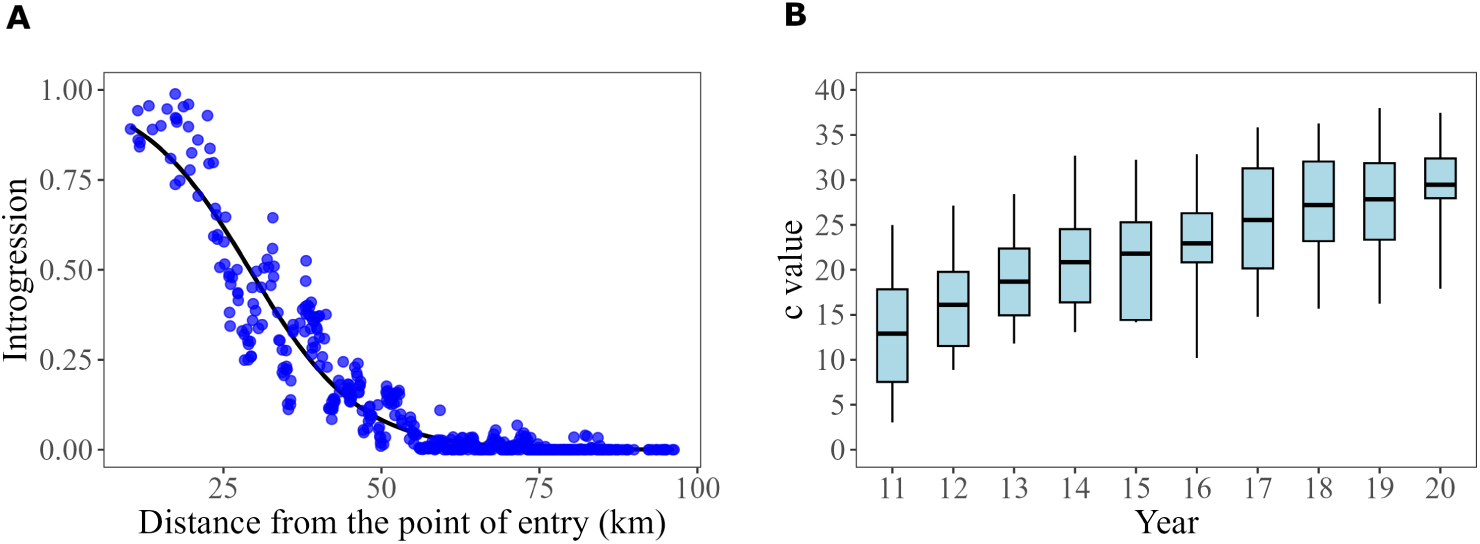
A) Scatter plot of the introgression levels in honeybee queens in relation to the distance from the central point of entry after 10 years of importation. Each blue dot represents a honeybee queen. The black line indicates the fitted cline to the scatter plot. B) Box plot of the annual changes in the *c* value (centre of the cline), across 10 years of import over replicates.

### 3.4 Halt of imports

To evaluate the ability of the population to recover from imports, we extended the spatially-aware scenario (point of entry) by halting imports in year 20 and we continued the simulation for another 10 years.

By year 20, there was a clear increase in the level of introgression, as shown in the results above for the spatially-aware scenario, reaching a value of 0.14 and a rate of introgression of 0.006/year. In the following 10 years without import, we observed a slow decay for the rate of introgression (−0.001/year) compared to the increase observed in the importation period. This led to a gradual decline in levels of introgression, decreasing to 0.13 by the end of the simulation (Figure 8). There was however a substantial variation between replicates on how these levels changed over time.

**Fig. 8.**
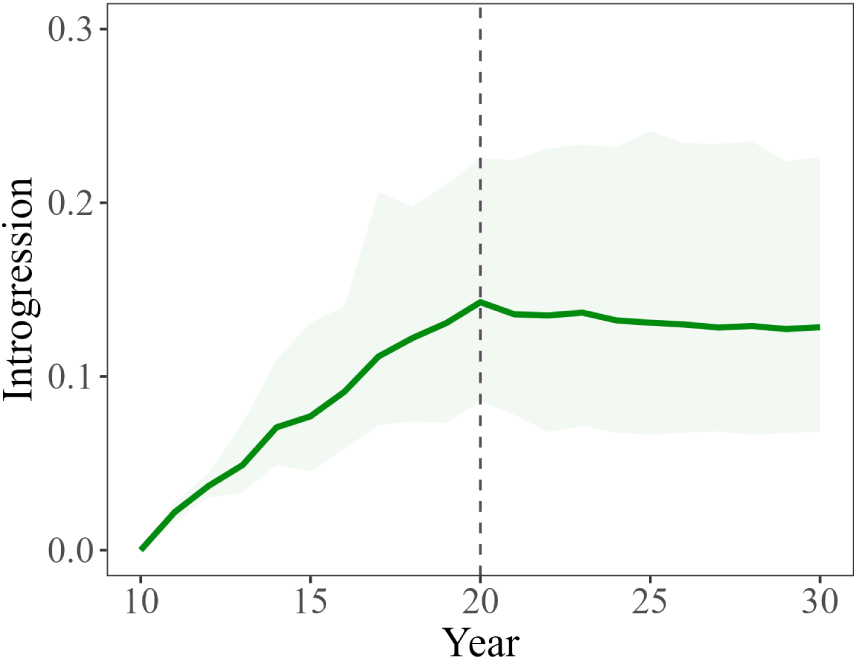
Trend in the mean level of introgression from year 10 to year 30 across replicates with corresponding shaded regions indicating the 95% quantiles. The dashed line marks the halt of import.

## 4 Discussion

In this study, we used a stochastic simulation model to quantify how the rate and extent of imports of non-native honey bee queens affects introgression, honey yield, and fitness in a native population, using Ireland as a case study. We showed that introgression rate and extent increased with the proportion of imported individuals, as expected. We also demonstrated the importance of genetic correlation of fitness across environments for the rate and spread of introgression and revealed the need for its accurate estimation. Lastly, we showed that the halt of importation alone does not lead to a reduction in the proportion of introgressed material, suggesting that once established, introgressed alleles can persist and continue to spread within populations. Our findings highlight the need for early and proactive conservation strategies of honey bees. The developed framework also provides tools and guidance for exploring hybridisation dynamics under varying ecological and genetic scenarios for other species.

### 4.1 Percentage of imports

Our simulated import rates captured the European spectrum: at the upper end, the UK imported 21,405 queens in 2020 (on the order of 10% of imports) (Department for Environment, Food and Rural Affairs 2021) while other European countries are reporting high introgression rates (Oleksa et al. 2011; Yanbaev 2024); mid-range values align with Ireland’s 4% in 2012, with higher figures during the COVID period (McCann and McCormack 2024); and at the lower end, several Nordic conservation areas restrict non-native introductions (e.g. Læsø in Denmark; protected zones in Nor-way), implying low import rates (Groeneveld et al. 2020). In Ireland, the reported introgression of 18-25% (Valentine et al. 2024; Smith et al. 2023) aligns with our simulation with 4% of imports that generated 18.3% of introgression after 10 years. Although expected, the consistent response and the agreement with observed data suggests that the model responds reliably to variation in key parameters and produces realistic results. This supports its suitability for informing conservation and management decisions. Furthermore, the results confirmed that the imports benefit genetic gain while disrupting mean fitness. Although expected, due to the simulated directional selection for honey yield in the non-native population, quantification of the response to increasing imports contributes to our understanding of population and phenotype dynamics.

### 4.2 Genetic correlation between traits in the two environments

Decreasing the genetic correlation for fitness had limited effect on the rate and magnitude of introgression. This was unexpected, but could be explained with the simulated natural selection pressure being too weak to purge non-native alleles. We assumed the native population to be at its fitness optimum and under stabilising selection, but also that fitness is a highly complex polygenic trait with low heritability. Taken together, these assumptions meant that introgressed non-native alleles were largely hidden to selection and could “silently” increase in frequency over time without major impacts on fitness. We hypothesise that simulating directional selection for fitness in the native population would result in a more efficient purging of non-native alleles.

The results also revealed that when genetic correlations between environments are low or negative, the introduction of non-native honeybees may reduce locally adapted fitness, despite short-term productivity gains. This is expected since a negative genetic correlation between two environments implies that the animals performing best in one environment perform worse in the other environment (e.g. Falconer 1952). Although GxE interactions have not been extensively studied in honeybees, some evidence suggests higher survival in their native environment compared to non-native environments (Ndiaye 2017; Meixner et al. 2015), and that importing non-adapted stock could decrease fitness and increase colony losses (Büchler et al. 2014; Henriques et al. 2018). Importantly, negative consequences may not be immediately apparent due to the polygenic nature of fitness, but could still have long-term consequences for local adaptation. In salmonids, introgression from domesticated or non-local stocks is associated with reduced fitness and altered life-history across wild populations, and modelling studies tracking fitness under different introgression pressures found that sustained introgression leads to clear declines in wild salmonids fitness (Castellani et al. 2018).

Our findings emphasize the need to estimate GxE in honeybees, a field severely understudied in all insects. The need to estimate and account for GxE in selection has already been recognised in livestock with the most advanced international systems being developed for cattle (Nilforooshan and Jorjani 2022). Understanding and quantifying how population’s performance and survival changes depending on the environment is crucial for managing imports, not only for researchers, but also for breeders and beekeepers making breeding decisions that shape the population. These groups may focus on immediate economic benefits while overlooking long-term consequences for population viability. This is crucial in regions such as Ireland, where the native subspecies *Apis mellifera mellifera* is under pressure from the importation of non-native honeybees (Hassett et al. 2018), and there is no governmental restriction of imports.

### 4.3 Point of entry

After 10 years of spatial simulation, introgression was detected in colonies located up to 5 times the mating range. While the general pattern of introgression spread is expected, quantifying the spatial extent and rate of spread provides useful insight into how quickly introgression can expand across landscapes.

Importantly, results highlight that even a single introduction point can lead to widespread introgression over relatively few generations (5–7 honeybee generations). By tweaking the simulation settings, the model can be used to assess spatial risk under different introduction scenarios, strategies aimed at limiting the spread of introgressed alleles, and design of conservation zones. Protected areas for *A. m. mellifera* in Nordic countries (Ruottinen et al. 2014) and the Læsø island conservation pro-gramme in Denmark (Pinto et al. 2014) demonstrate that geographic isolation can limit introgression. Where geographic isolation is not available, as is largely the case in Ireland, the results suggest that restricting import locations and building restriction zones around native colonies could slow, though not eliminate, introgression spread.

Our simulation is the first in honeybees incorporating the spatial structure that captures key processes governing gene flow across landscapes. This represents an important step towards more biologically realistic simulations, particularly when evaluating management strategies that depend on geographic constraints. Other simulation tools also recognised the need for a spatial component that enables a joint study of populations’ genetics and ecology (Chevy et al. 2024; Petr et al. 2023). Such modelling has already proven useful for managed pollinators around greenhouses or release hotspots and for large mammals with focal introductions, such as red–sika deer in mixed estates (Seabra et al. 2019; Goodman et al. 1999).

### 4.4 Halt of imports

Ten years after halting imports following a 10-year import period we observed only a slight reduction in introgression. We hypothesise this is due to the simulated selection pressure being too weak. Under stabilising selection on a low-heritable poly-genic fitness trait, introgressed alleles decline only slowly once established, as seen in other species (Hedrick 2009; Karlsson et al. 2016). For example, seven generations after releases ceased, two wild sea trout populations still retained large amounts of introgressed material (Bekkevold et al. 2025). Similarly, nine years after changes to rainbow-trout stocking in southeastern British Columbia, westslope cutthroat trout at open sites still carried substantial rainbow-trout ancestry (Bennett 2009). In managed populations, the promotion of non-native alleles is also exacerbated by artificial maintenance by farmers/beekeepers due to positive effects on production despite potential negative effects on fitness.

These results highlight the difficulty of reversing introgression once it has occurred, especially in species with high dispersal like honeybees (Tallmon et al. 2004), and a need for an early implementation of prevention measures. In Ireland, where introgression remains relatively low, measures such as limiting or banning non-native imports could avoid following the trajectory already observed in other countries (U.S. House of Representatives, Committee on Agriculture 1962).

### 4.5 Model assumptions and transferability of the results

Simulation studies are a valuable tool for testing population management decisions before practical implementation. To produce realistic outcomes, the simulation must use estimates of parameters from real population studies. When not available, we must make educated assumptions or test outcomes over a grid of the parameter space that we identify as most relevant based on the knowledge and experience from other systems. In this study, we decided to make assumptions for the parameters we either had some insight for or that tend to be non- or less variable in the nature, and vary across the parameters that are unknown or vary in the real-life as well.

We assumed directional selection for honey-yield in the non-native population, which is realistic since honey yield is improved through artificial selection that applies truncation selection. For the fitness trait in the evaluation phase, we assumed that both populations are at the optimal value for fitness and under stabilising selection around this value. While this is a reasonable assumption (e.g. Kingsolver et al. (2001)), other modes of selection, such as directional or disruptive, could be applied. We also assumed random mating in a non-spatial or spatial context, and did not account for mating preference or reproductive barriers between subspecies. However, there is evidence of behavioural differences between honey bees that would result in assortative mating, such as different vertical position of drones at mating (Koeniger et al. 1989), that would decrease the rate of introgression.

On the other hand, we varied the extent of import and GxE interaction for the fitness trait. Varying over some parameters is an opportunity to explore outcomes that might pertain to different settings and assess the sensibility of the outcome to certain parameters in the same setting. However, it can also appear as a limitation when implementing population management strategies for a specific population. Although previous studies provide qualitative insights into the impact of non-native honeybees on native populations (Büchler et al. 2014; Meixner et al. 2015; Ndiaye 2017; Henriques et al. 2018), there is still limited quantitative estimates. In particular, knowledge of GxE interaction remains scarce, despite its central role in determining the consequences of introgression. This limitation extends to other systems, including well studied organisms such as *Drosophila*, where the environmental factors shaping hybrid performance and the long-term stability of introgressed alleles remain poorly understood (Miller and Matute 2017). Moreover, there is a lack of quantified extent and distribution of dominance effects. The latter could confer hybrid vigour, which could facilitate introgression by increasing the fitness or production of hybrids. Demonstrating the sensibility of the results, especially to the extent of GxE interaction, emphasizes the need for accurate estimation and quantification of population and trait parameters.

The developed model and assessment of parameter impact can also be used to model other social species or guide the simulations for even more distantly related taxa. SIMplyBee is built on AlphaSimR (Gaynor et al. 2021), which is intended to simulate non-social livestock and plant populations and breeding programs. SIMply-Bee thus retains all of AlphaSimR’s general functionality, including using an external coalescence simulator ((Chen et al. 2009)) for genome simulation. While SIMply-Bee specifies parameters for honeybee genome and demography, users can specify any demographic model, as well as mutation and recombination rates. Furthermore, although SIMplyBee was built to accommodate peculiarities of social organisms, it can also simulate simpler scenarios such as monoandrous mating or organisms without social structure. In the latter, the code essentially boils down to AlphaSimR’s code, meaning the same principle and code flow can be used to model other species (e.g (Obšteter et al. 2023a),(Fritsche-Neto et al. 2023)).

We observed high variability across replicates indicating a huge amount of randomness in introgression outcomes driven by the randomness of genetic inheritance, coupled with selection on phenotypes that are a noisy representation of genetic value of individuals. Due to required computational resources, we simulated only 500 colonies and a surface of 150*km*^2^. Although this represents a fraction of the Irish population, the simulated population dynamics and results still hold if import is limited to a few entry points with limited migration outside of these areas. However, a smaller population might have exacerbated introgression and weakened the selection pressure on the imported individuals due to a sizeable drift effect. Future work could focus on increasing the number of the simulation and replicates to better quantify the outcomes and the variability itself.

As shown in our simulation, even a small percentage of introductions can lead to introgression of non-native alleles whose effects in a new environment are unknown. In many natural systems experiencing biological invasions or secondary contact (Rhymer and Simberloff 1996), detailed records of admixture rates are rarely available, making simulation approaches like ours particularly valuable for studying introgression dynamics. Our findings suggest that similar processes may operate wherever human-mediated movement brings divergently adapted lineages into contact (Muhlfeld et al. 2009; Castellani et al. 2018; Bekkevold et al. 2025).

### 4.6 Application to the management of honeybee populations

Assuming significant differences in production between native and non-native populations, we observed that introgression will rapidly spread from the point of entry, increasing genetic value for honey yield and decreasing genetic value for fitness. So the pertinent question is: “What is the way forward”? Although import can increase production and commercial interest in the short-term, it generates a long-term decrease in the fitness of the native population. While beekeepers might overcome this with management, beekeeping practices should be developed alongside native populations and their requirements. Additionally, the increase in commercially relevant traits may have a negative impact on the fitness of free-living honeybees which are not under human management and therefore cannot benefit from such mitigation strategies. In the long term, introgression will depend on the complex interplay of a number of factors, including benefits in production and costs of adjusting the management interest of beekeepers that want imports and those that do not, selective breeding actions, knowledge about the native and non-native populations and systems, and strength and direction of natural selection. Climate change will also likely be a significant factor in this process, as changing environmental conditions may alter selective pressures, the relative performance of native and non-native genotypes, and increase the likelihood of secondary contact between previously isolated populations.

The best course of action to prevent loosing the identity of the native populations and their adaptations due to imports is “conservation-by-use”. Some commercial beekeepers import non-native honeybees to improve short-term productivity and profitability. However, while there is no evidence that non-native honeybees perform better in Ireland than native bees, both anecdotal and scientific studies have reported adaptations of *A. m. mellifera* to the climate in Ireland, including morphological (Valentine et al. 2024; Ruttner et al. 1990) and behavioural adaptations (Cooper and Denwood 1986). If this is the case, these adaptations would have evolved over many years and could be impacted or even lost due to introgression. By establishing selective breeding programmes aimed at enhancing production and workability traits of native populations under local environmental conditions, beekeepers could improve commercial viability while preserving the past genetic adaptations of the populations. In fact, countries known for exporting their honeybee subspecies globally have established selective breeding programmes. For example, Italy has been selectively breeding for higher productivity and docility for decades (Bar-Cohen et al. 1978), while Germany, Poland, and Switzerland, have been successful at breeding for increased brood viability or hygienic behavior (Uzunov et al. 2014).

## 5 Conclusion

Our work offers useful guidelines for modelling introgression in honeybees as well as other species with similar dispersal patterns. Here, we provide an open-source stochastic simulation and evaluation model with focus on complex traits, which can be revised and extended as needed and when new knowledge becomes available.

Our results demonstrate that although importing non-native honeybees can increase production traits, it also increases introgression of the non-native alleles, which is associated with a decline in fitness and the loss of native adaptation to the environment. Moreover, the spread of non-native alleles can be fast and substantial with continued imports and introgression cannot be reversed quickly, resulting in long-term integration of the introgressed alleles. These findings highlight the importance of strategies to manage genetically distinct and locally adapted subspecies of honeybees. Imposing strict import regulations may be a necessary solution to preserve native populations as has been applied by some countries and regions. However, it is important to understand and address the reasons for importing non-native honeybees. A comprehensive approach would be (1) to restrict levels/locations of imports of non-native honeybees, (2) to enhance economically important traits of the native honeybees via selective breeding, (3) to raise awareness of the expected level of production of different genotypes in different environments, and (4) to communicate about the risk of short- versus long-term effects of introgressing non-native alleles. To support such an approach, it is important to expand population and quantitative genetics research of honeybee populations.

## 6 Acknowledgments

IC and GM acknowledge that the project was funded by the Department of Agriculture, Food and the Marine under RFT 234005 - for the provision of research services regarding the Native Irish Honey bee (*Apis mellifera mellifera*).

LS and GG acknowledge the support from BBSRC ISP grants (BBS/E/D/30002275, BBS/E/RL/230001A, and BBS/E/RL/230001C) and support from the BBSRC DTP (EASTBio) CASE PhD studentship with AbacusBio.

JO acknowledges the core financing of the Slovenian Research and Innovation Agency (programme P4-0133 “Sustainable agriculture”).

## 7 Author information

### 7.1 Authors’ contributions

IC: conceptualisation, simulation coding, data curation, formal analysis, visualisation, writing – original draft.

LS: conceptualisation, simulation coding, methodology, writing – review and editing.

GM: conceptualisation, funding acquisition, supervision, project administration, writing – review and editing.

GG: conceptualisation, funding acquisition, supervision, project administration, methodology, validation, writing – review and editing.

JO: conceptualisation, supervision, simulation coding, methodology, validation, writ-ing – review and editing.

### 7.2 Corresponding author

Correspondence to Jana Obsteter & Irene de Carlos.

## 8 Availability of data and material

The datasets generated and/or analysed during the current study are available in the idecarlos_honeybee_import repository, https://github.com/HighlanderLab/idecarlos_honeybee_import.

## 9 Ethics declarations

### 9.2 Research Ethics statement

Research ethics committee’s approval was not required for this study.

## 10 Supplementary information

Link to the GitHub repository with code for the simulations: https://github.com/HighlanderLab/idecarlos_honeybee_import.

**Additional file 1.** ANOVA results for introgression and genetic values under different import and correlation scenarios.

|  | Rate of<br>introgression | Introgression<br>at year 20 | Honey yield<br>at year 20 | Fitness<br>at year 20 |
| --- | --- | --- | --- | --- |
| Percentage of imports | <2e-16 | <2e-16 | 7.69e-15 | 3.99e-05 |
| Fitness genetic correlations | 0.4760 | 0.2210 | 0.2260 | 0.0093 |

ANOVA p-values for the rate of introgression, introgression at year 20, honey yield at year 20, and fitness at year 20 across different import percentages and genetic correlation scenarios.

**Additional file 2.**
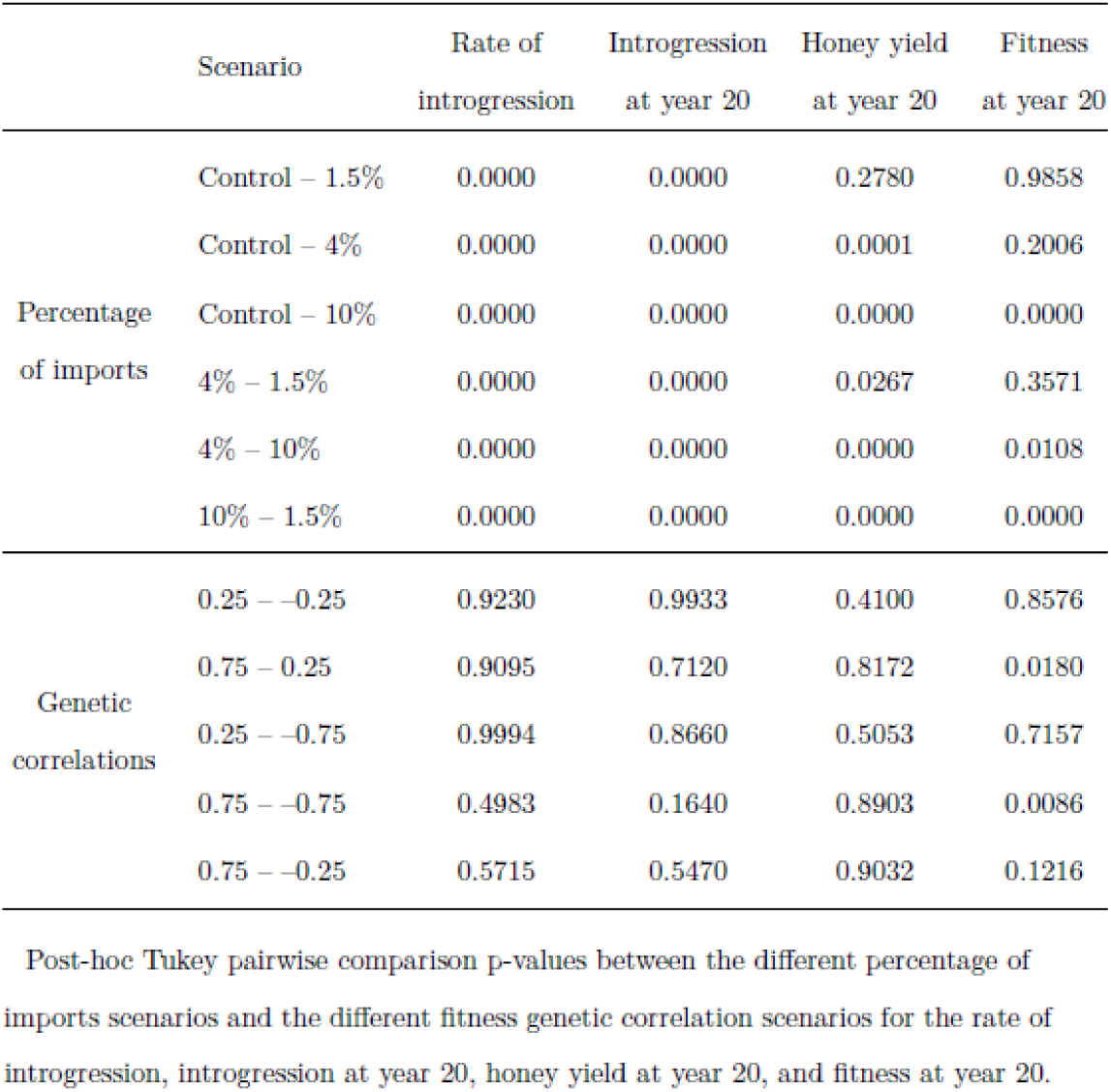
Tukey post-hoc comparisons.

**Additional file 3.**
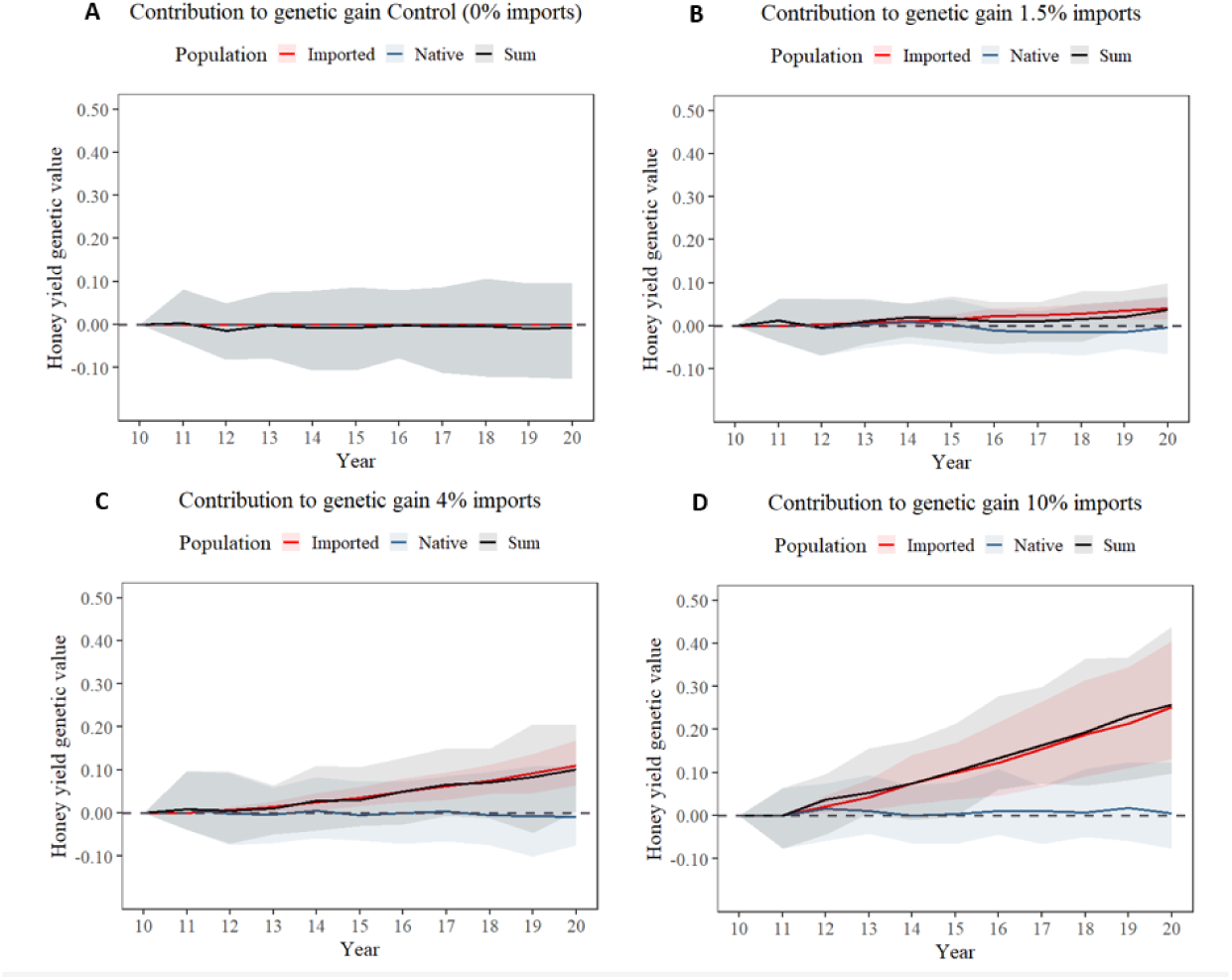
Contribution to genetic gain of honey yield from year 10 to year 20, comparing the imported (red line) and native contributions (blue line) to the overall honey yield genetic value for the Irish population or Sum (black line). This is represented across varying percentages of imports of C-lineage bees to the Irish population: A) Control (0% imports), B) 1.5% imports, C) 4% imports, and D) 10% imports. Shaded regions indicated the 95% confidence intervals. The contribution of imported C-lineage honeybees increased the genetic value of honey yield for the Irish population. This increase became more evident as the percentage of imports was incremented.

**Additional file 4.**
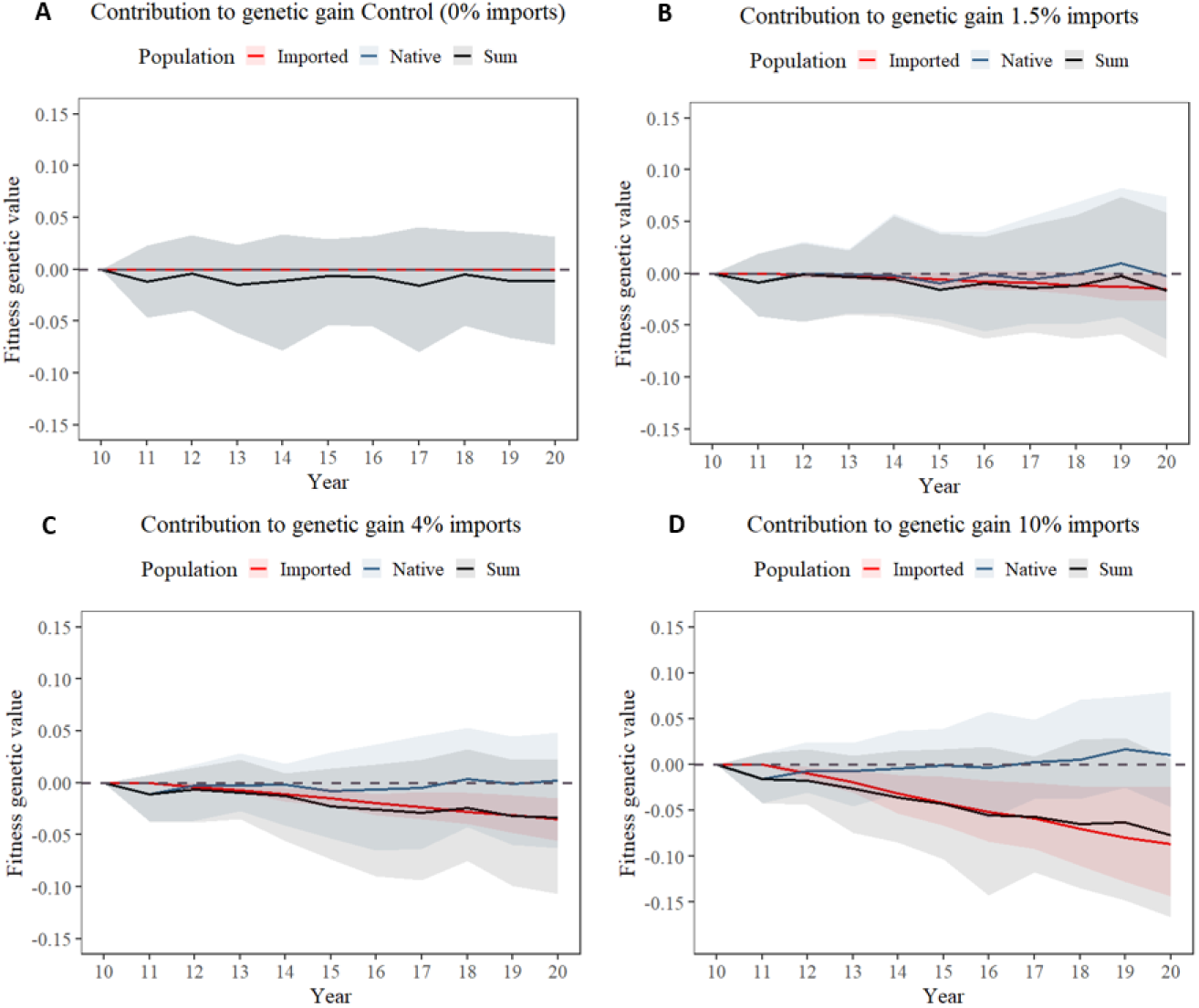
Contribution to genetic gain of fitness from year 10 to year 20, comparing the imported (red line) and native contributions (blue line) to the overall fitness genetic value for the Irish population or Sum (black line). This is represented across varying percentages of imports of C-lineage bees to the Irish population: A) Control (0% imports), B) 1.5% imports, C) 4% imports, and D) 10% imports. Shaded regions indicated the 95% confidence intervals. The contribution of imported C-lineage honeybees decreased the genetic value of fitness for the Irish population. This decrease became more evident as the percentage of imports was incremented.

**Additional file 5.**
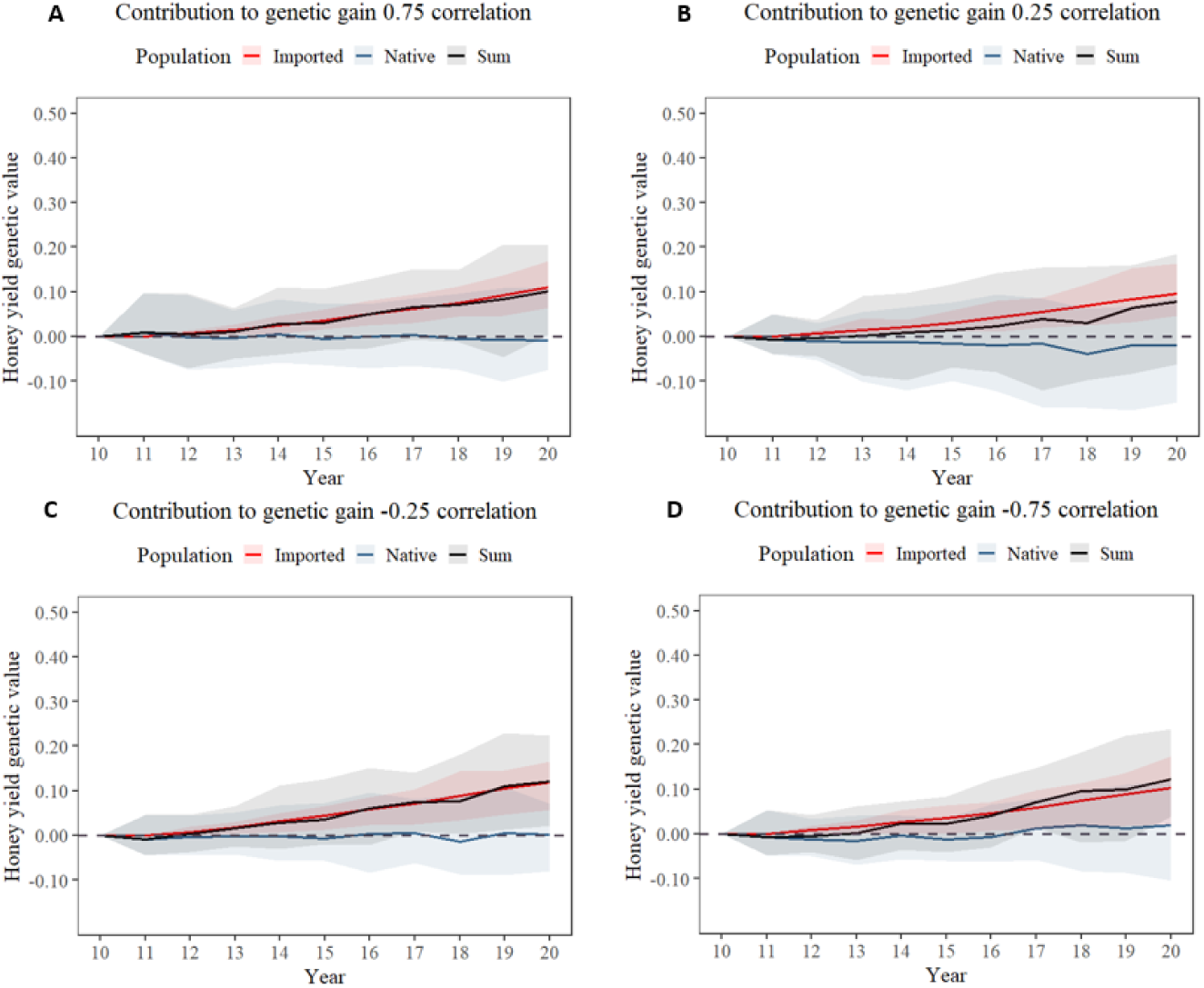
Contribution to genetic gain of honey yield from year 10 to year 20, comparing the imported (red line) and native contributions (blue line) to the overall honey yield genetic value for the Irish population or Sum (black line). This is represented across varying genetic correlations between fitness traits: A) Genetic correlation of 0.75, B) Genetic correlation of 0.25, C) Genetic correlation of −0.25, and D) Genetic correlation of −0.75. Shaded regions indicated the 95% confidence intervals. The contribution of imported C-lineage honeybees increased the genetic value of honey yield for the Irish population. However there was no clear pattern among the different fitness genetic correlations.

**Additional file 6.**
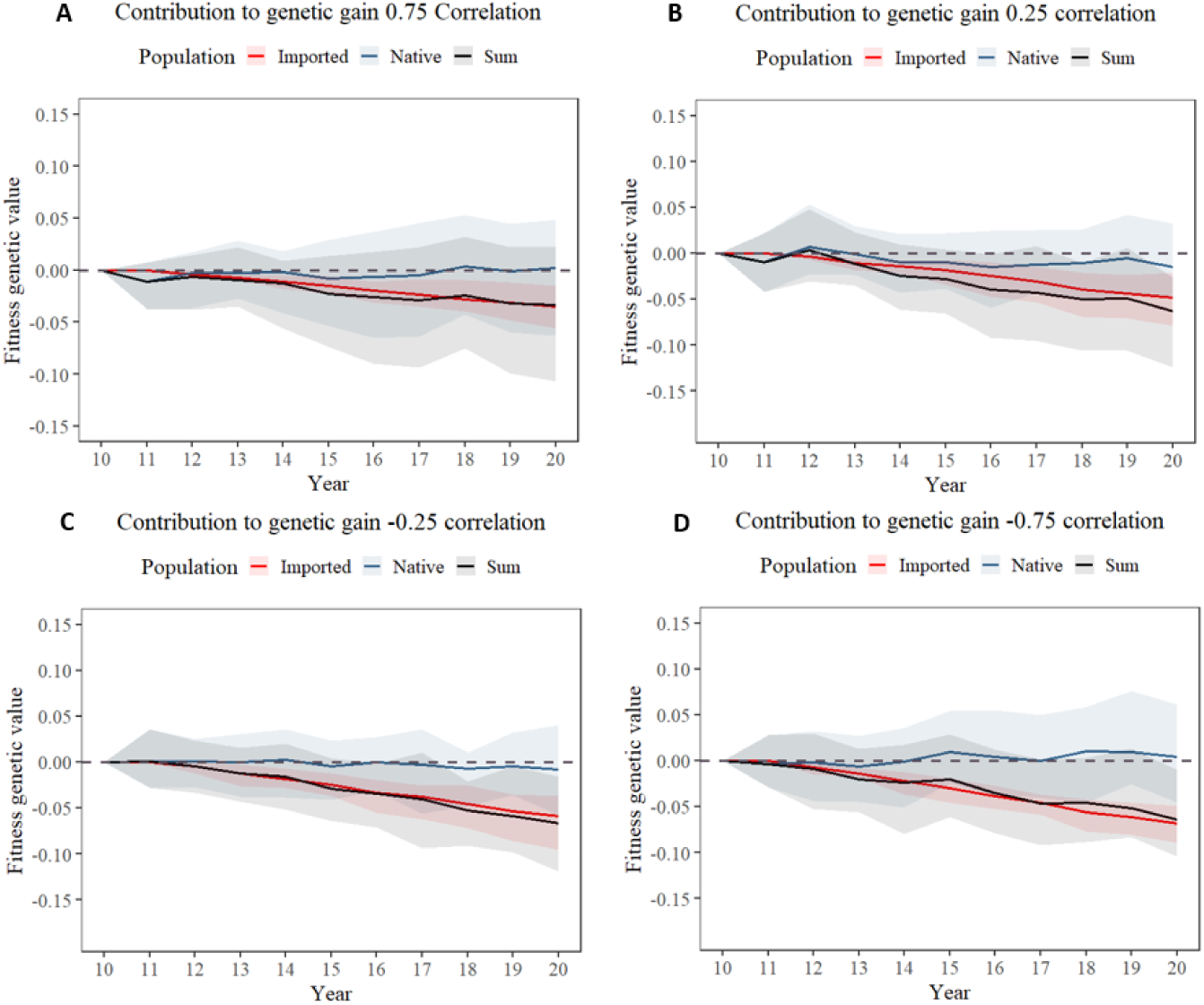
Contribution to genetic gain of fitness from year 10 to year 20, comparing the imported (red line) and native contributions (blue line) to the overall fitness genetic value for the Irish population or Sum (black line). This is represented across varying genetic correlations between fitness traits: A) Genetic correlation of 0.75, B) Genetic correlation of 0.25, C) Genetic correlation of −0.25, and D) Genetic correlation of −0.75. Shaded regions indicated the 95% confidence intervals. The contribution of the imported C-lineage honeybees is in generally negative. The contribution became increasingly negative as the fitness genetic correlation was decreased.

**Additional file 7.**
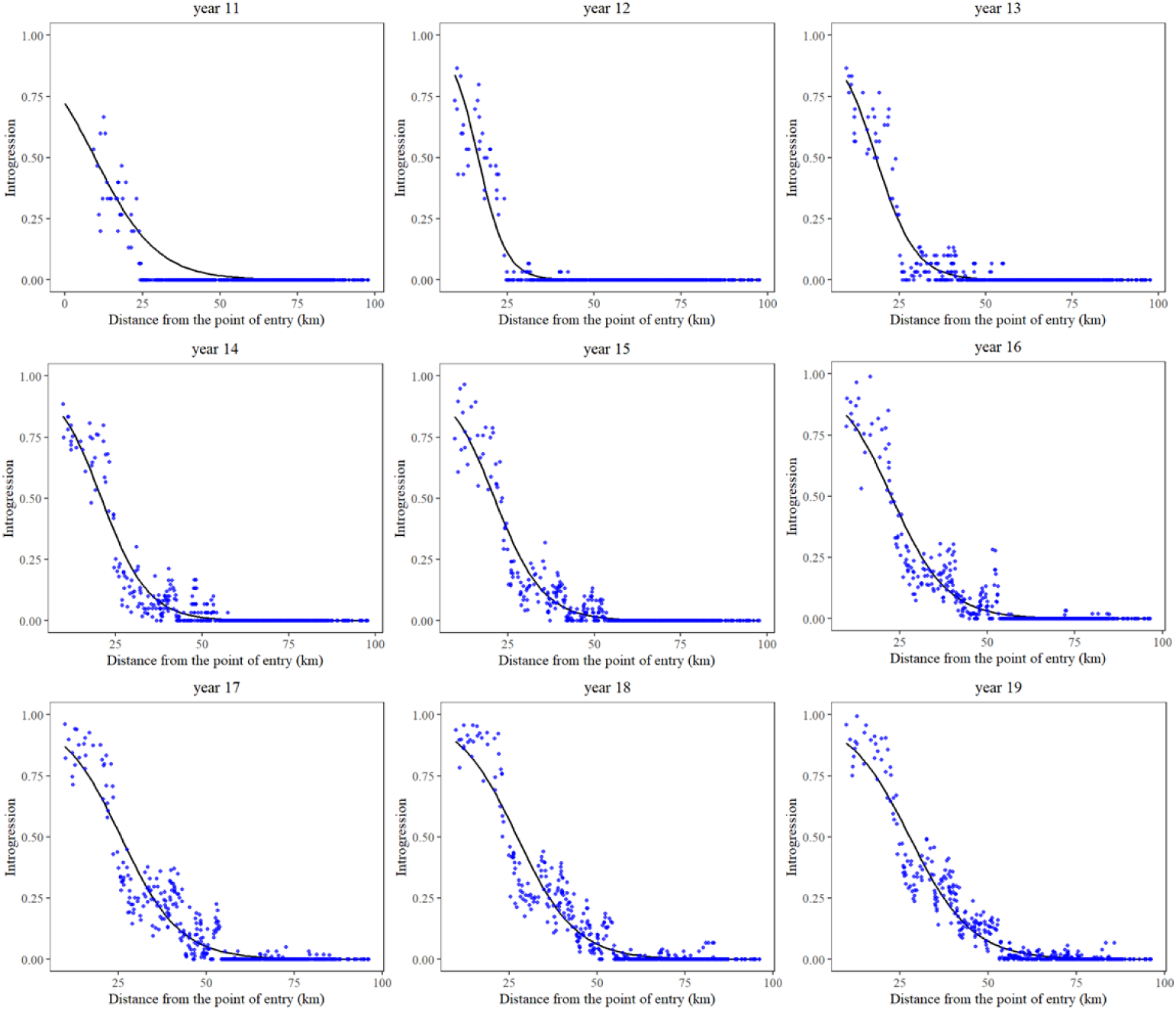
Scatter plot of the introgression levels in honeybee colonies across the distance from the central point of entry on the first (year 11) to the 9th (year 19) year of importation. Each blue dot corresponds to a honeybee colony, with the y-axis indicating the level of introgression and the x-axis representing the distance from the point of entry, which is marked at 0 km. The black line indicates the fitted cline to the scatter plot.

**Additional file 8.**
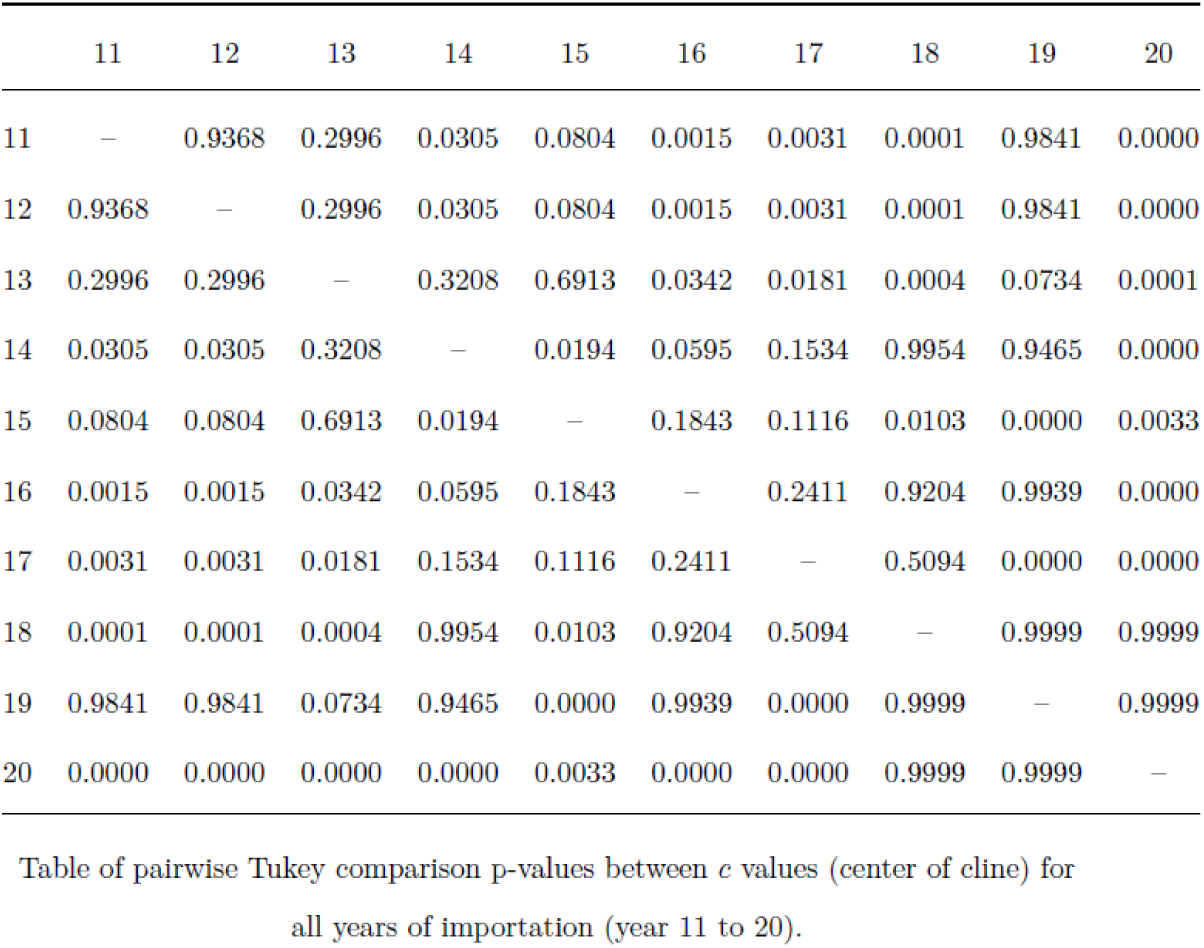
Tukey p-value matrix for cline centers pairwise comparisons.

|  | 11 | 12 | 13 | 14 | 15 | 16 | 17 | 18 | 19 | 20 |
| --- | --- | --- | --- | --- | --- | --- | --- | --- | --- | --- |
| 11 | – | 0.9368 | 0.2996 | 0.0305 | 0.0804 | 0.0015 | 0.0031 | 0.0001 | 0.9841 | 0.0000 |
| 12 | 0.9368 | – | 0.2996 | 0.0305 | 0.0804 | 0.0015 | 0.0031 | 0.0001 | 0.9841 | 0.0000 |
| 13 | 0.2996 | 0.2996 | – | 0.3208 | 0.6913 | 0.0342 | 0.0181 | 0.0004 | 0.0734 | 0.0001 |
| 14 | 0.0305 | 0.0305 | 0.3208 | – | 0.0194 | 0.0595 | 0.1534 | 0.9954 | 0.9465 | 0.0000 |
| 15 | 0.0804 | 0.0804 | 0.6913 | 0.0194 | – | 0.1843 | 0.1116 | 0.0103 | 0.0000 | 0.0033 |
| 16 | 0.0015 | 0.0015 | 0.0342 | 0.0595 | 0.1843 | – | 0.2411 | 0.9204 | 0.9939 | 0.0000 |
| 17 | 0.0031 | 0.0031 | 0.0181 | 0.1534 | 0.1116 | 0.2411 | – | 0.5094 | 0.0000 | 0.0000 |
| 18 | 0.0001 | 0.0001 | 0.0004 | 0.9954 | 0.0103 | 0.9204 | 0.5094 | – | 0.9999 | 0.9999 |
| 19 | 0.9841 | 0.9841 | 0.0734 | 0.9465 | 0.0000 | 0.9939 | 0.0000 | 0.9999 | – | 0.9999 |
| 20 | 0.0000 | 0.0000 | 0.0000 | 0.0000 | 0.0033 | 0.0000 | 0.0000 | 0.9999 | 0.9999 | – |

Table of pairwise Tukey comparison p-values between *c* values (center of cline) for all years of importation (year 11 to 20).

## References

Allendorf FW, Leary RF, Spruell P, Wenburg JK. The problem with species: Reincarnation or sacrilege? Hybridization and the conservation of species. Trends in Ecology & Evolution. 2001;16(11):613–622. URL https://doi.org/10.1016/S0169-5347(01)02290-X, doi: 10.1016/S0169-5347(01)02290-X.

Bančič J, Greenspoon P, Gaynor CR, Gorjanc G. Plant breeding simulations with AlphaSimR. Crop Science. 2024;65(1). URL http://dx.doi.org/10.1002/csc2.21312, doi: 10.1002/csc2.21312.

Bar-Cohen R, Alpern G, Bar-Anan R. Progeny testing and selecting Italian queens for brood area and honey production. Apidologie. 1978;9(2):95–100.

Bekkevold D, Glover KA, Jimenez-Mena B, Besnier F, Nielsen EE. Genetic recovery of two wild seatrout populations following long-term stocking with non-native conspecifics. Molecular Ecology. 2025;34:e70036. Open Access, doi: 10.1111/mec.70036.

Benjamin MM, Cheyenne P, Quinn L, Daniel LP, Yaniv B, Molly S. The genomic consequences of hybridization. eLife. 2021;10:e69016. URL 10.7554/eLife.69016.

Bennett SN. Levels of introgression in westslope cutthroat trout populations: Nine years after changes to rainbow trout stocking programs in southeastern British Columbia. North American Journal of Fisheries Management. 2009;29(5):1271–1282. doi: 10.1577/M08-048.1.

Beye M, Gattermeier I, Hasselmann M, Gempe T, Schioett M, Baines JF, et al. Exceptionally high levels of recombination across the honey bee genome. Genome Res. 2006 Nov;16(11):1339–1344. doi: 10.1101/gr.5680406.

Bleeker W, Martin KG. Interspecific hybridisation between alien and native plant species in Germany and its consequences for native biodiversity. Biological Conservation. 2007;137(2):248–253. doi: 10.1016/j.biocon.2007.02.004.

Brodschneider R, Gray A, Adjlane N, Ballis A, Brusbardis V, Charrière JD, et al. Multicountry loss rates of honeybee colonies during winter 2016/2017 from the COLOSS survey. Journal of Apicultural Research. 2018 May;57(3):452–457. URL https://doi.org/10.1080/00218839.2018.1460911, publisher: Taylor & Francis _eprint: https://doi.org/10.1080/00218839.2018.1460911, doi: 10.1080/00218839.2018.1460911.

Büchler R, Costa C, Hatjina F, Andonov S, Meixner MD, Conte YL, et al. The influence of genetic origin and its interaction with environmental effects on the survival of Apis mellifera L. colonies in Europe. Journal of Apicultural Research. 2014 Jan;53(2):205–214. URL https://doi.org/10.3896/IBRA.1.53.2.03, publisher: Taylor & Francis _eprint: https://doi.org/10.3896/IBRA.1.53.2.03, doi: 10.3896/IBRA.1.53.2.03.

Castellani M, Heino M, Gilbey J, Araki H, Svåsand T, Glover KA. Modeling fit-ness changes in wild Atlantic salmon populations faced by spawning intrusion of domesticated escapees. Evolutionary Applications. 2018;11(6):1010–1025. URL https://doi.org/10.1111/eva.12615, doi: 10.1111/eva.12615.

Chen GK, Marjoram P, Wall JD. Fast and flexible simulation of DNA sequence data. Genome Res. 2009;19(1):136–142. doi: 10.1101/gr.083634.108.

Chevy ET, Min J, Caudill V, Champer SE, Haller BC, Rehmann CT, et al. Population genetics meets ecology: a guide to individual-based simulations in continuous landscapes. bioRxiv. 2024;URL http://dx.doi.org/10.1101/2024.07.24.604988, doi: 10.1101/2024.07.24.604988.

Cooper BA, Denwood P. The Honeybees of the British Isles. Codnor, Derby: British Isles Bee Breeders’ Association; 1986.

Corrêa AS, Cordeiro EM, Omoto C. Agricultural insect hybridization and implications for pest management. Pest Management Science. 2019;75(11):2857–2864. URL https://scijournals.onlinelibrary.wiley.com/doi/abs/10.1002/ps.5495, doi: 10.1002/ps.5495. https://scijournals.onlinelibrary.wiley.com/doi/pdf/10.1002/ps.5495.

Department for Environment, Food and Rural Affairs. Bee importation: In 2020 we imported over 21,000 queens to Great Britain. UK Government; 2021.

Department of Agriculture, Food and the Marine. Beekeepers Census 2019. Dublin, Ireland: Government of Ireland; 2019.

Department of Agriculture, Food and the Marine. Annual honey bee queen import data for Ireland. Government of Ireland; 2024.

Derryberry EP, Derryberry GE, Maley JM, Brumfield RT. hzar: hybrid zone analysis using an R software package. Molecular Ecology Resources. 2014;14(3):652–663. URL https://onlinelibrary.wiley.com/doi/abs/10.1111/1755-0998.12209, doi: 10.1111/1755-0998.12209. https://onlinelibrary.wiley.com/doi/pdf/10.1111/1755-0998.12209.

Falconer DS. The Problem of Environment and Selection. The American Naturalist. 1952 sep;86(830):293–298. URL https://www.jstor.org/stable/2457811.

Fritsche-Neto R, Ali J, De Asis EJ, Allahgholipour M, Labroo MR. Improving hybrid rice breeding programs via stochastic simulations: number of parents, number of hybrids, tester update, and genomic prediction of hybrid performance. Theoretical and Applied Genetics. 2023;137(1):3. doi: 10.1007/s00122-023-04508-6.

Gaynor RC, Gorjanc G, Hickey JM. AlphaSimR: An R-package for Breeding Pro-gram Simulations. G3: Genes|Genomes|Genetics. 2021;11. doi: 10.1093/g3journal/jkaa017.

Glenny W, Cavigli I, Daughenbaugh KF, Radford R, Kegley SE, Flenniken ML. Honeybee (Apis mellifera) colony health and pathogen composition in migra-tory beekeeping operations involved in California almond pollination. PLOS ONE. 2017 08;12:1–24. URL https://doi.org/10.1371/journal.pone.0182814, doi: 10.1371/journal.pone.0182814.

Goodman SJ, Barton NH, Swanson G, Abernethy K, Pemberton JM. Introgression through rare hybridization: A genetic study of a hybrid zone between red and sika deer (genus Cervus) in Argyll, Scotland. Genetics. 1999;152(1):355–371. URL https://doi.org/10.1093/genetics/152.1.355, doi: 10.1093/genetics/152.1.355.

Gray A, Adjlane N, Arab A, Ballis A, Brusbardis V, Bugeja Douglas A, et al. Honeybee colony loss rates in 37 countries using the COLOSS sur-vey for winter 2019–2020: the combined effects of operation size, migration and queen replacement. Journal of Apicultural Research. 2023 Mar;62(2):204–210. URL https://doi.org/10.1080/00218839.2022.2113329, publisher: Taylor & Francis _eprint: https://doi.org/10.1080/00218839.2022.2113329, doi: 10.1080/00218839.2022.2113329.

Groeneveld LF, Kirkerud LA, Dahle B, Sunding M, Flobakk M, Kjos M, et al. Conservation of the dark bee (Apis mellifera mellifera): Estimating C-lineage introgression in Nordic breeding stocks. Acta Agriculturae Scandinavica, Section A – Animal Science. 2020;69(3):157–168. URL https://doi.org/10.1080/09064702.2020.1770327, doi: 10.1080/09064702.2020.1770327.

Guichard M, Phocas F, Neuditschko M, Basso B, Dainat B. An Overview of Selection Concepts Applied to Honey Bees. Bee World. 2023;100(1):2–8. doi: 10.1080/0005772X.2022.2147702.

Guichard M, Phocas F, Neuditschko M, Basso B, Dainat B. An Overview of Selection Concepts Applied to Honey Bees. Bee World. 2023 Jan;100(1):2–8. URL https://doi.org/10.1080/0005772X.2022.2147702, publisher: Taylor & Francis _eprint: https://doi.org/10.1080/0005772X.2022.2147702, doi: 10.1080/0005772X.2022.2147702.

Hassett J, Browne KA, McCormack GP, Moore E, Society NIHB, Soland G, et al. A significant pure population of the dark European honey bee (Apis mellifera mellifera) remains in Ireland. Journal of Apicultural Research. 2018 May;57(3):337–350. URL https://doi.org/10.1080/00218839.2018.1433949, publisher: Taylor & Francis _eprint: https://doi.org/10.1080/00218839.2018.1433949, doi: 10.1080/00218839.2018.1433949.

Hatjina F, Costa C, Ralph B, Uzunov A, Dražić M, Filipi J, et al. Population dynamics of European honey bee genotypes under different environmental conditions. Journal of Apicultural Research. 2014 May;53:233–247. doi: 10.3896/IBRA.1.53.2.05.

Havill NP, Davis G, Mausel DL, Klein J, McDonald R, Jones C, et al. Hybridization between a native and introduced predator of Adelgidae: An unintended result of classical biological control. Biological Control. 2012 Dec;63(3):359–369. URL http://dx.doi.org/10.1016/j.biocontrol.2012.08.001, doi: 10.1016/j.biocontrol.2012.08.001.

Hedrick PW. Conservation Genetics and North American Bison (Bison bison). Journal of Heredity. 2009 Jul;100(4):411–420. URL https://doi.org/10.1093/jhered/esp024, doi: 10.1093/jhered/esp024.

Henriques D, Browne KA, Barnett MW, Parejo M, Kryger P, Freeman TC, et al. High sample throughput genotyping for estimating C-lineage introgression in the dark honeybee: an accurate and cost-effective SNP-based tool. Scientific Reports. 2018 Jun;8(1):8552. URL https://www.nature.com/articles/s41598-018-26932-1, number: 1 Publisher: Nature Publishing Group, doi: 10.1038/s41598-018-26932-1.

Hoppe A, Du M, Bernstein R, Tiesler FK, Kärcher M, Bienefeld K. Substantial Genetic Progress in the International Apis mellifera carnica Population Since the Implementation of Genetic Evaluation. Insects. 2020;11(11):768. URL 10.3390/insects11110768.

Huxel GR. Rapid displacement of native species by invasive species: effects of hybridization. Biological Conservation. 1999 Jan;89(2):143–152. doi: 10.1016/S0006-3207(98)00153-0.

Jensen AB, Palmer KA, Chaline N, Raine NE, Tofilski A, Martin SJ, et al. Quantifying honey bee mating range and isolation in semi-isolated valleys by DNA microsatellite paternity analysis. Conservation Genetics. 2005 Jul;6(4):527–537. URL https://doi.org/10.1007/s10592-005-9007-7, doi: 10.1007/s10592-005-9007-7.

Jones B, Semmence N. Imported Honey Bees: Risks and Mitigation. BBKA News Incorporating The British Bee Journal. 2021 Sep;68:295–296. Published by Fera Science Ltd and APHA.

Karlsson S, Diserud OH, Fiske P, Hindar K. Widespread genetic introgression of escaped farmed Atlantic salmon in wild salmon populations. ICES Journal of Marine Science. 2016;73(10):2488–2498. URL https://doi.org/10.1093/icesjms/fsw121, doi: 10.1093/icesjms/fsw121.

Kingsolver JG, Hoekstra HE, Hoekstra JM, Berrigan D, Vignieri SN, Hill CE, et al. The Strength of Phenotypic Selection in Natural Populations. The American Naturalist. 2001;157(3):245–261. doi: 10.1086/319193.

Koeniger G, Koeniger N, Pechhacker H, Ruttner F, Berg S. Assortative mating in a mixed population of European Honeybees, Apis mellifera ligustica and Apis mellifera carnica. Insectes Sociaux. 1989 Jun;36(2):129–138. URL https://doi.org/10.1007/BF02225908, doi: 10.1007/BF02225908.

Kruuk LE, Clutton-Brock TH, Slate J, Pemberton JM, Brotherstone S, Guinness FE. Heritability of fitness in a wild mammal population. Proceedings of the National Academy of Sciences of the United States of America. 2000 Jan;97(2):698–703. doi: 10.1073/pnas.97.2.698.

Lewontin RC, Birch LC. Hybridization as a source of variation for adaptation to new environments. Evolution. 1966;20(3):315–336. doi: 10.2307/2406633.

Mallet J. Hybridization as an invasion of the genome. Trends in Ecology & Evolution. 2005;20(5):229–237. URL https://doi.org/10.1016/j.tree.2005.02.010, doi: 10.1016/j.tree.2005.02.010.

Martínez-López V, Ruiz C, De la Rúa P. “Migratory beekeeping and its influence on the prevalence and dispersal of pathogens to managed and wild bees”. International Journal for Parasitology: Parasites and Wildlife. 2022;18:184–193. URL https://www.sciencedirect.com/science/article/pii/S2213224422000499, doi: 10.1016/j.ijppaw.2022.05.004.

Mattila HR, Otis GW. Influence of Pollen Diet in Spring on Development of Honey Bee (Hymenoptera: Apidae) Colonies. Journal of Economic Entomology. 2006 Jun;99(3):604–613. URL https://doi.org/10.1093/jee/99.3.604, doi: 10.1093/jee/99.3.604.

Maucourt S, Fortin F, Robert C, Giovenazzo P. Genetic Parameters of Honey Bee Colonies Traits in a Canadian Selection Program. Insects. 2020 Sep;11(9):587. doi: 10.3390/insects11090587.

McCann M, McCormack GP. Increased levels of introgression evident in Irish honey bees. Journal of Apicultural Research. 2024;63(1):205–207. URL 10.1080/00218839.2023.2262872.

McKinney ML, Lockwood JL. Biotic homogenization: a few winners replacing many losers in the next mass extinction. Trends in ecology and evolution. 1999;14(11):450–453. URL 10.1016/S0169-5347(99)01679-1.

Meixner MD, Kryger P, Costa C. Effects of genotype, environment, and their interactions on honey bee health in Europe. Current Opinion in Insect Science. 2015;10:177–184. URL https://www.sciencedirect.com/science/article/pii/S2214574515000899, social Insects * Vectors and Medical and Veterinary Entomology, doi: 10.1016/j.cois.2015.05.010.

Metcalf JL, Siegle MR, Martin AP. Hybridization dynamics between Colorado’s native cutthroat trout and introduced rainbow trout. Journal of Heredity. 2008;99(2):149–156. URL https://doi.org/10.1093/jhered/esm118, doi: 10.1093/jhered/esm118.

Miller CJJ, Matute DR. The Effect of Temperature on Drosophila Hybrid Fitness. G3: Genes Genomes Genetics. 2017;7(2):377–385. URL https://doi.org/10.1534/g3.116.034926, doi: 10.1534/g3.116.034926.

Minozzi G, Lazzari B, De Iorio MG, Costa C, Carpana E, Crepaldi P, et al. Whole-Genome Sequence Analysis of Italian Honeybees Apis mellifera. Animals (Basel). 2021;11(5):1311. URL 10.3390/ani11051311.

Muhlfeld CC, Kalinowski ST, McMahon TE, Taper ML, Painter S, Leary RF, et al. Hybridization rapidly reduces fitness of a native trout in the wild. Biology Letters. 2009;5(3):328–331. URL https://doi.org/10.1098/rsbl.2009.0033, doi: 10.1098/rsbl.2009.0033.

Ndiaye N, Ndiaye N, editor.: Analysis of Iberian honey bee (Apis mellifera iberiensis) colony phenotypes across two different origins in Portugal: Searching for evidence of genotype by environment interactions, Mestrado em Recursos Florestais. Instituto Politécnico de Bragança; 2017. URL https://bibliotecadigital.ipb.pt/handle/10198/14431.

Nilforooshan M, Jorjani H. Invited review: A quarter of a century—International genetic evaluation of dairy sires using MACE methodology. Journal of Dairy Science. 2022;105(1):3–21. doi: 10.3168/jds.2021-21104.

Obšteter J, Holl J, Hickey JM, Gorjanc G. AlphaPart—R implementation of the method for partitioning genetic trends. Genetics Selection Evolution. 2021 Mar;53(1):30. URL https://doi.org/10.1186/s12711-021-00600-x, doi: 10.1186/s12711-021-00600-x.

Obšteter J, Jenko J, Pocrnic I, Gorjanc G. Investigating the benefits and perils of importing genetic material in small cattle breeding programs via simulation. Journal of Dairy Science. 2023;106(8):5593–5605. doi: 10.3168/jds.2022-23132.

Obšteter J, Strachan LK, Bubnič J, Prešern J, Gorjanc G. SIMplyBee: an R package to simulate honeybee populations and breeding programs. Genetics, Selection, Evolution : GSE. 2023 May;55:31. URL https://www.ncbi.nlm.nih.gov/pmc/articles/PMC10169377/, doi: 10.1186/s12711-023-00798-y.

Oleksa A, Chybicki I, Tofilski A, Burczyk J. Nuclear and mitochondrial patterns of introgression into native dark bees (Apis mellifera mellifera) in Poland. JOURNAL OF APICULTURAL RESEARCH. 2011;50(2):116–129. doi: 10.3896/IBRA.1.50.2.03.

Petr M, Haller BC, Ralph PL, Racimo F. slendr: a framework for spatiotemporal population genomic simulations on geographic landscapes. Peer Community Journal. 2023;3. URL https://peercommunityjournal.org/articles/10.24072/pcjournal.354/, doi: 10.24072/pcjournal.354.

Pinto AM, Henriques D, Chavez-Galarza J, Kryger P, Garnery L, van der Zee R, et al. Genetic integrity of the Dark European honey bee (Apis mellifera mellifera) from protected populations: a genome-wide assessment using SNPs and mtDNA sequence data. JOURNAL OF APICULTURAL RESEARCH. 2014;53(2, SI):269–278. doi: 10.3896/IBRA.1.53.2.08.

Pook T, Schlather M, Simianer H. MoBPS - Modular Breeding Program Simulator. G3 Genes|Genomes|Genetics. 2020;10(6):1915–1918. URL https://doi.org/10.1534/g3.120.401193, doi: 10.1534/g3.120.401193.

Péntek-Zakar E, Oleksa A, Borowik T, Kusza S. Population structure of honey bees in the Carpathian Basin (Hungary) confirms introgression from surrounding subspecies. Ecol Evol. 2015;5(23):5456–5467. URL 10.1002/ece3.1781.

Quilodran CS, Austerlitz F, Currat M, Montoya-Burgos JI. Cryptic Bio-logical Invasions: a General Model of Hybridization. Scientific Reports. 2018;8(1):2414. URL https://www.nature.com/articles/s41598-018-20543-6, doi: 10.1038/s41598-018-20543-6.

R Core Team. R: A Language and Environment for Statistical Computing. Vienna, Austria: R Foundation for Statistical Computing; 2024.

Rangel J, Keller J, Tarpy D. The effects of honey bee (Apis mellifera L.) queen reproductive potential on colony growth. Insectes Sociaux. 2013 Feb;60. doi: 10.1007/s00040-012-0267-1.

Rhymer JM, Simberloff D. Extinction by Hybridization and Introgression. Annual Review of Ecology and Systematics. 1996;27(1):83–109. URL https://doi.org/10.1146/annurev.ecolsys.27.1.83, _eprint: https://doi.org/10.1146/annurev.ecolsys.27.1.83, doi: 10.1146/annurev.ecolsys.27.1.83.

De la Rua P, Jaffé R, Dall’Olio R, Muñoz I, Serrano J. Biodiversity, conservation and current threats to European honeybees. Molecular Ecology. 2009;40(3):263–284. URL 10.1051/apido/2009027.

Ruottinen L, Peer B, Kantanen J, Kristensen T, Kettunen A. Status and Conservation of the Nordic Brown Bee: Final Report. Nord Gen. 2014;15.

Ruttner F, Milner E, Dews JE. The Dark European Honeybee. British Isles Bee Breeders’ Association; 1990.

Schultz KM, Passino KM, Seeley TD. The mechanism of flight guidance in honeybee swarms: subtle guides or streaker bees? Journal of Experimental Biology. 2008 Oct;211(20):3287–3295. URL https://doi.org/10.1242/jeb.018994, doi: 10.1242/jeb.018994.

Seabra SG, Silva SE, Nunes VL, Sousa VC, Martins J, Marabuto E, et al. Genomic signatures of introgression between commercial and native bumblebees, Bombus terrestris, in western Iberian Peninsula—Implications for conservation and trade regulation. Evolutionary Applications. 2019;12(6):1218–1232. URL https://doi.org/10.1111/eva.12732, doi: 10.1111/eva.12732.

Seitz N, Traynor KS, Steinhauer N, Rennich K, Wilson ME, Ellis JD, et al. A national survey of managed honey bee 2014–2015 annual colony losses in the USA. Journal of Apicultural Research. 2015 Aug;54(4):292–304. URL http://www.scopus.com/inward/record.url?scp=84969850493&partnerID=8YFLogxK, doi: 10.1080/00218839.2016.1153294.

Smith S, Moro A, McCormack GP. Exploring a Potential Avenue for Beekeeping in Ireland: Safeguarding Locally Adapted Honeybees for Breeding Varroa-Resistant Lines. Insects. 2023 Oct;14(10):827. URL https://www.ncbi.nlm.nih.gov/pmc/articles/PMC10607453/, doi: 10.3390/insects14100827.

Tallmon DA, Luikart G, Waples RS. The alluring simplicity and complex reality of genetic rescue. Trends in Ecology & Evolution. 2004 Sep;19(9):489–496. doi: 10.1016/j.tree.2004.07.003.

Thomas DS. Honeybee Ecology. Princeton University Press; 2016. URL https://press.princeton.edu/books/hardcover/9780691639352/honeybee-ecology, iSBN: 9780691639352.

U S House of Representatives, Committee on Agriculture. Importation of Honey Bees: Report to Accompany H.R. 8050. Washington, D.C.: 87th Congress, 2nd Session; 1962.

Uzunov A, Costa C, Panasiuk B, Meixner M, Kryger P, Hatjina F, et al. Swarming, defensive and hygienic behaviour in honey bee colonies of different genetic origin in a pan-European experiment. Journal of Apicultural Research. 2014 05;53:248–260. doi: 10.3896/IBRA.1.53.2.06.

Valentine A, Moro A, Briggs E, Collier B, Sandoval K, Binetti C, et al. Introgressive hybridisation puts the distinctive population of Apis mellifera mellifera in Ireland at risk: evidence from a multidisciplinary approach. Journal of Apicultural Research. 2024 10;64:1–15. doi: 10.1080/00218839.2024.2404297.

Wallberg A, Han F, Wellhagen G, Dahle B, Kawata M, Haddad N, et al. A worldwide survey of genome sequence variation provides insight into the evolutionary history of the honeybee Apis mellifera. Nature Genetics. 2014 Oct;46(10):1081–1088. doi: 10.1038/ng.3077.

Yanbaev Y. Spatial analysis of genetic variation in a natural population of the dark forest bee (Apis mellifera mellifera L.) from the Southern Urals, Russia. International Journal of Environmental Studies. 2024;URL https://doi.org/10.1080/00207233.2022.2058768, doi: 10.1080/00207233.2022.2058768.

Yang S, Wang L, Huang J, Zhang X, Yuan Y, Chen JQ, et al. Parent-progeny sequencing indicates higher mutation rates in heterozygotes. Nature. 2015;523(7561):463–467. doi: 10.1038/nature14649.

